# Astrocytes render memory flexible

**DOI:** 10.1101/2021.03.25.436945

**Authors:** Wuhyun Koh, Mijeong Park, Ye Eun Chun, Jaekwang Lee, Hyun Soo Shim, Mingu Gordon Park, Sunpil Kim, Hyunji Kang, Soo-Jin Oh, Junsung Woo, Heejung Chun, Seungeun Lee, Jinpyo Hong, Jiesi Feng, Yulong Li, Hoon Ryu, Jeiwon Cho, C. Justin Lee

## Abstract

Cognitive flexibility is an essential ability to adapt to changing environment and circumstances. NMDAR has long been implicated in cognitive flexibility, but the precise molecular and cellular mechanism is not well understood. Here, we report that astrocytes regulate NMDAR tone through Best1-mediated glutamate and D-serine release, which is critical for cognitive flexibility. Co-release of D-serine and glutamate is required for not only homosynaptic LTD but also heterosynaptic LTD, which is induced at unstimulated synapses upon release of norepinephrine and activation of astrocytic α1-AR during homosynaptic LTP. Remarkably, heterosynaptic LTD at unstimulated synapses during memory acquisition is required for later repotentiation LTP during reversal learning, laying a foundation for flexible memory and cognitive flexibility. Our study sheds light on the pivotal role of astrocytes in orchestrating multiple synapses during memory formation and determining the fate of consolidated memory to be retained as a flexible memory.

**Highlights:**

1. Astrocytes regulate NMDAR tone via Best1-mediated glutamate and D-serine release
2. Activation of astrocytic α1-AR induces heterosynaptic LTD via NMDAR tone
3. Heterosynaptic LTD is required for repotentiation LTP and spatial reversal learning
4. Astrocytic regulation of NMDAR tone is critical for metaplasticity and flexible memory

## Introduction

The flexibility of memory is as important as the formation of memory because an environment and circumstances are not static, but dynamically changing. When necessary, previously acquired memories should be flexibly adjusted to adapt to the changing environment. This ability is generally termed as cognitive flexibility (Tello-Ramos et al., 2019). Cognitive flexibility has been reported to decline in several diseases, for instance, autism spectrum disorder (ASD) (D’Cruz et al., 2013), schizophrenia (Wobrock et al., 2009), and early stages of Alzheimer’s disease (AD) (Etienne et al., 2013; Guarino et al., 2018), in which a hypofunction of N-methyl-D-aspartate receptor (NMDAR) is implicated (Gandal et al., 2012; Huang et al., 2012; Lee and Zhou, 2019). However, little is known about how NMDAR hypofunction affects cognitive flexibility. In the hippocampus, NMDAR-dependent long-term depression (LTD) is proposed to be associated with spatial reversal learning (Duffy et al., 2008; Li et al., 2015; Morice et al., 2007; Nicholls et al., 2008), a hippocampus-dependent form of cognitive flexibility (Izquierdo et al., 2017). Of the synaptic NMDAR (synNMDAR) and extrasynaptic NMDAR (exNMDAR) classified according to location, the latter has been suggested to be particularly important for LTD in the hippocampus (Liu et al., 2013; Lu et al., 2001). The current mediated by exNMDAR is also referred to as “tonic NMDAR” or “NMDAR tone” due to its slow-time-scale or continuous occupancy/activation of exNMDAR by an ambient glutamate (Le Meur et al., 2007) and co-agonists (i.e. D-serine and glycine). Although exNMDAR appears to be crucial for LTD and cognitive flexibility, it has not been yet clear which cell regulates NMDAR tone for exNMDAR currents in the brain.

D-serine, one of the co-agonists that could constitute NMDAR tone, has been extensively investigated in hippocampal LTD and spatial reversal learning. For example, exogenous D-serine was shown to augment hippocampal LTD (Duffy et al., 2008; Zhang et al., 2008) and spatial reversal learning (Duffy et al., 2008). In addition, a transgenic mouse with a loss-of-function mutation in D-amino acid oxidase (DAAO), a key catabolic enzyme for D-serine, showed enhanced spatial reversal learning (Labrie et al., 2009). In contrast, a depletion of endogenous D-serine by DAAO was shown to impair and reduce hippocampal LTD (Papouin et al., 2012; Zhang et al., 2008). While these results suggest that D-serine-mediated NMDAR tone can facilitate spatial reversal learning, the precise molecular and cellular mechanism of how endogenous D-serine is regulated and facilitates spatial reversal learning is not well understood. Importantly, it has been still unclear and controversial whether the cellular source of D-serine is astrocytes (Papouin et al., 2017) or neurons (Wolosker et al., 2017).

Recently, optogenetic activation of channelrhodopsin-2 (ChR2) in astrocytes has been shown to induce an increase in hippocampal NMDAR tone (Shen et al., 2017) and hippocampal LTD (Navarrete et al., 2019). This increase in NMDAR tone was reduced by the treatment of NPPB (Shen et al., 2017), which blocks anion channels including the Ca^2+^-activated, glutamate-permeable anion channel Best1 (Oh and Lee, 2017), suggesting that glutamate released through Best1 from astrocytes (Park et al., 2013; Woo et al., 2012) contributes to hippocampal NMDAR tone and LTD induction. However, in place of exogenous ChR2, which induces hippocampal LTD, an endogenous molecule that activates astrocytes has not been identified yet. Previously, it has been reported that chemogenetic activation of locus coeruleus (LC) can restore spatial reversal learning in early stages of AD model (Rorabaugh et al., 2017). Interestingly, norepinephrine (NE), synthesized in LC neurons, is known to induce hippocampal LTD through α1-adrenergic receptor (α1-AR) (Dyer-Reaves et al., 2019; Scheiderer et al., 2004), which is predominantly localized in astrocytes (Hertz et al., 2010). It has been later demonstrated that astrocytes respond to NE through α1-AR by an increase in cytosolic Ca^2+^ (Ding et al., 2013), which can possibly open Best1 in Ca^2+^-dependent manner. However, it is not known whether NE, through activation of astrocytic α1-AR, can cause an increase in NMDAR tone and hippocampal LTD via Best1.

Activation of astrocytes was shown to induce not only homosynaptic LTD (Navarrete et al., 2019), but also heterosynaptic LTD (Chen et al., 2013a), which was initially documented to occur at the unstimulated synapses accompanying homosynaptic long-term potentiation (LTP) at the stimulated synapses (Lynch et al., 1977; Scanziani et al., 1996). In early times, homosynaptic LTD induced by low-frequency stimulation (LFS) has been thought to be associated with spatial reversal learning (Nicholls et al., 2008). In contrast, heterosynaptic LTD has been suggested to enable metaplasticity (Chen et al., 2013b), which is defined as a plasticity of synaptic plasticity (Abraham and Bear, 1996; Hulme et al., 2014). However, it is not known whether heterosynaptic LTD is associated with spatial reversal learning. Paradoxically, it has been also demonstrated that spatial reversal learning is either enhanced in mice with decreased homosynaptic LTD (Zhang and Wang, 2013) or diminished in mice with increased homosynaptic LTD (Rutten et al., 2011). These findings challenge the initial notion that homosynaptic LTD is responsible for spatial reversal learning and raise the possibility of heterosynaptic LTD to be more closely linked to spatial reversal learning. Nevertheless, it has not been studied whether heterosynaptic LTD through activation of astrocytes is associated with spatial reversal learning.

In the present study, we have investigated how astrocytes regulate heterosynaptic LTD and metaplasticity, and thereby contribute to spatial reversal learning. We employed cell-type specific genetic manipulations, *ex vivo* electrophysiological recordings, sniffer patch recordings, cutting-edge biosensor for NE, and behavioral assays to investigate whether astrocytes can regulate NMDAR tone by releasing D-Serine and glutamate. Subsequently, we further investigated the role for NMDAR tone in the heterosynaptic LTD, metaplasticity and cognitive flexibility. We found that CA1-hippocampal astrocytes indeed dynamically control heterosynaptic LTD during an induction of homosynaptic LTP through NMDAR tone regulation via Best1. Furthermore, we found that this heterosynaptic LTD becomes a basis for the flexible memory which is required for spatial reversal learning and cognitive flexibility.

## Results

### Astrocytes regulate hippocampal NMDAR tone through Best1

It has been suggested that astrocytic Ca^2+^ is an important signaling molecule for the release of gliotransmitters (Sahlender et al., 2014; Semyanov et al., 2020), including glutamate (Lee et al., 2007; Parpura and Haydon, 2000) and D-serine (Henneberger et al., 2010; Takata et al., 2011). Thus, we hypothesized that if hippocampal astrocytes regulate NMDAR tone in a Ca^2+^-dependent manner, an inhibition of astrocytic Ca^2+^ would lead to reduced NMDAR tone. To investigate whether astrocytes regulate NMDAR tone in hippocampus, exNMDAR current was measured in CA1 pyramidal neuron, while Ca^2+^ signaling in nearby astrocytes is suppressed by Ca^2+^-clamping. To measure exNMDAR current, a shift in the whole-cell current in response to a treatment of 50 μM (2R)-amino-5-phosphonovaleric acid (APV), the NMDAR blocker, was quantified under voltage-clamping at +40mV in the presence of other inhibitors (20 μM CNQX for AMPAR, 10 μM Bicuculline for GABA_A_R, 10 μM CGP 55845 for GABA_B_R, 10 μM Strychnine for GlyR) (Figure 1A). To inhibit astrocytic Ca^2+^ signaling, BAPTA, a Ca^2+^ chelator, and Alexa Fluor 488, a fluorescent indicator, were loaded into astrocytes through a patch pipette (Figures 1B and S1A), as previously described (Kwak et al., 2020; Serrano et al., 2006; Shigetomi et al., 2008). We observed that exNMDAR current was significantly reduced when astrocytes were loaded with BAPTA (+BAPTA), compared to without BAPTA (-BAPTA) condition (Figures 1C and 1D). These results indicate that astrocytes regulate hippocampal NMDAR tone in a Ca^2+^-dependent manner.

**Figure 1.**
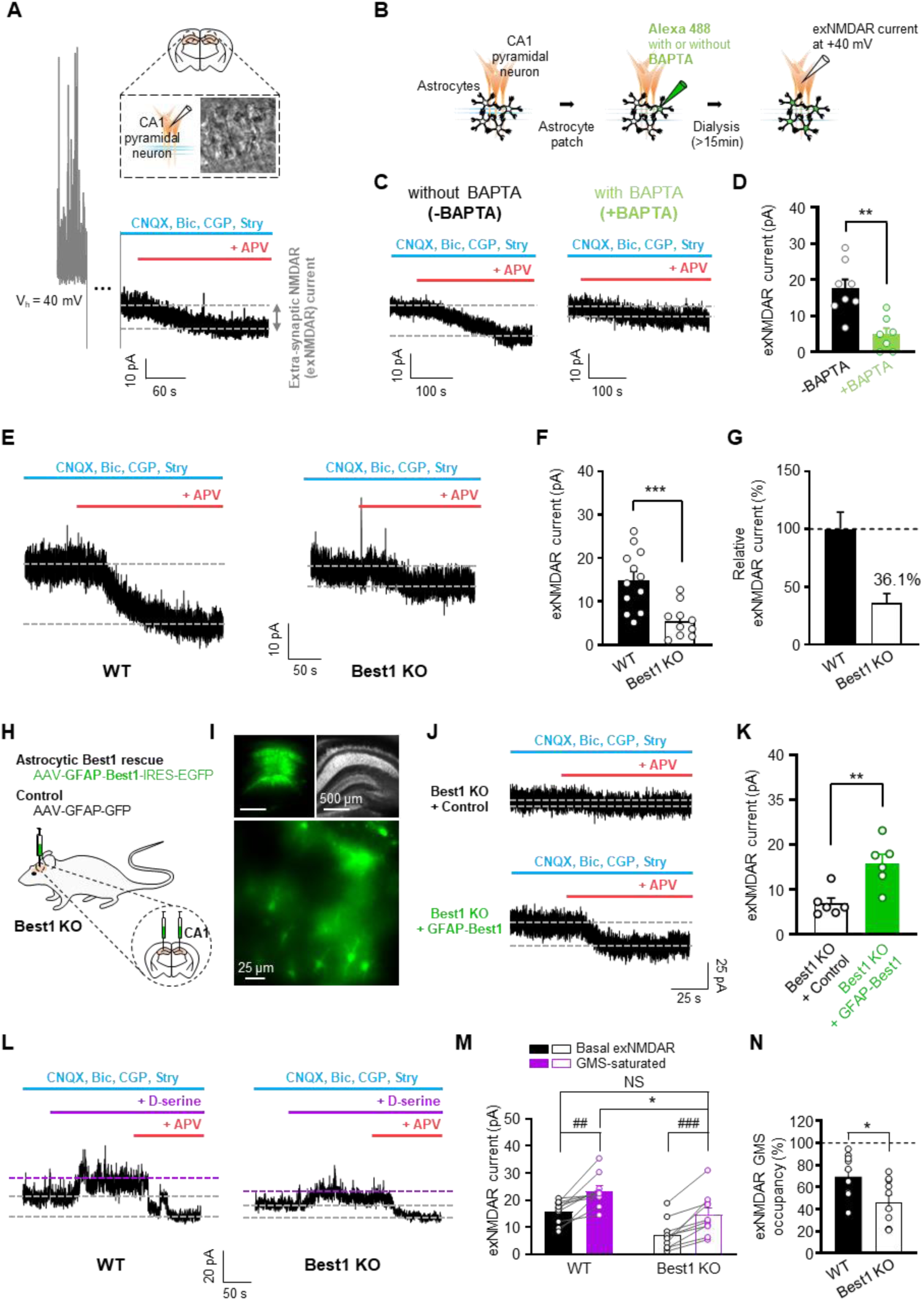
Astrocytes regulate hippocampal NMDAR tone through Best1. (A) Extrasynaptic NMDAR (exNMDAR) current recording in hippocampal CA1 pyramidal neuron. To isolate NMDAR-mediated current, CNQX, Bicuculline (Bic.), CGP55845 (CGP.), and Strychnine (Stry.) were used. APV-sensitive current in voltage holding at +40mV. (B) Astrocytic Ca^2+^ chelation with BAPTA dialysis. (C) Representative traces of exNMDAR current with or without BAPTA dialysis. (D) Summary graph of exNMDAR current with or without BAPTA dialysis. (E-G) Representative traces of exNMDAR current (E), summary graph of exNMDAR current (F), and relative exNMDAR current to WT (G) in WT and Best1 KO. (H-K) Scheme of astrocytic Best1 rescue in CA1 of Best1 KO (H), images showing virus expression (I), representative traces of exNMDAR current (J) and summary graph of exNMDAR current in each condition (K). (L) Application of 100 µM D-serine during exNMDAR measurement. (M) Summary graph of exNMDAR current before (black) and after D-serine treatment (purple) in each condition. (N) Estimated exNMDAR GMS occupancy (%). Individual dots refer to cells. Data are represented as mean ± SEM. *p < 0.05; **p < 0.01; ***p < 0.001; Mann Whitney test (D and K) or unpaired-t test (F, M, N). #p < 0.05; ##p < 0.01; ###p < 0.001; paired t-test (M).

It has been previously reported that Bafilomycin A treatment did not affect exNMDAR current, indicating that vesicular release does not majorly contribute to exNMDAR current (Le Meur et al., 2007). Based on this finding, we hypothesized that a non-vesicular, Ca^2+^-dependent channel-mediated mechanism might be involved, and tested the possibility that Best1-mediated gliotransmission contributes to exNMDAR current using Best1 knockout (KO) mice (Marmorstein et al., 2006). We found that exNMDAR current was significantly decreased in Best1 KO, compared to the wild type (WT) mice (Figures 1E-1F). This decrease in exNMDAR current in Best1 KO was on average 63.9% of that in WT (Figure 1G). To test if the decreased exNMDAR current in Best1 KO is due to the lack of Best1-mediated gliotransmission in astrocytes, we over-expressed *Best1* full-clone under GFAP promoter in hippocampal astrocytes of Best1 KO by injecting AAV-GFAP-Best1-IRES-EGFP virus into CA1 hippocampus (Figures 1H, 1I and S1B). We found that the decreased exNMDAR current in Best1 KO was significantly recovered in astrocyte-specific Best1 rescue group (GFAP-Best1), but not in control group (GFAP-GFP) (Figures 1J and 1K), indicating that the decreased exNMDAR current in Best1 KO is caused by reduced Best1-mediated gliotransmission from astrocytes. The decreased exNMDAR current was not due to a decreased protein expression of NMDAR, as evidenced by consistent protein expression level of GluN1, an indispensable subunit for functional NMDAR (Figure S2). These results suggest that astrocytic Ca^2+^-activated anion channel Best1 mediates NMDAR tone in hippocampus.

NMDAR tone can be attributed to both glutamate and NMDAR co-agonist (i.e. D-serine or glycine), as both are required for NMDAR activation. To dissect the contribution of glutamate component to NMDAR tone, exNMDAR glycine modulatory site (GMS) was fully occupied by an exogenous application of 100μM D-serine (Figure 1L). The application of D-serine increased exNMDAR current in WT by 47.9% (before, 15.9±1.4 pA; after, 23.5±1.9 pA; mean ± SEM) and Best1 KO by 105% (before, 7.2±1.8 pA; after, 14.7±2.4 pA; mean ± SEM) (Figures 1L and 1M), implying that GMS was not fully occupied in both genotypes under basal condition. More importantly, we found that the GMS-saturated exNMDAR current was significantly reduced in Best1 KO to 62.6% (Figure 1M), indicating a reduced ambient level of glutamate in Best1 KO. Remarkably, we observed that when D-serine was applied to Best1 KO, the GMS-saturated exNMDAR current was significantly recovered to the level of WT (Figure 1M), indicating that D-serine supplement is sufficient to fully rescue the impaired NMDAR tone of Best1 KO to the WT level. To measure the ambient level of NMDAR co-agonist, GMS-occupancy of exNMDAR was calculated by dividing the basal exNMDAR current by GMS-saturated exNMDAR current for each genotype (Figures 1M and 1N). The percentage of GMS-occupancy of exNMDAR was significantly reduced in Best1 KO by 33.4% (WT, 69.4±6.0%; Best1 KO, 46.2±6.6%; mean ± SEM) (Figure 1N), indicating that Best1 KO has reduced NMDAR co-agonist as well as glutamate. Furthermore, we observed a similarly reduced GMS-occupancy of synNMDAR in Best1 KO as well as in *Best1* gene-silencing condition, which was fully rescued by an astrocyte-specific *Best1* rescue (Figures S3A-S3N). Taken together, these results indicate that astrocytes regulate NMDAR tone through Best1 by modulating the ambient level of both glutamate and co-agonist.

### D-serine and glutamate are co-released from astrocyte through Best1

To investigate which co-agonist, D-serine or glycine, is released from astrocytes in a Ca^2+^-dependent manner, we conducted a sniffer-patch experiment with a solitary primary cultured astrocyte and a sensor cell, as previously described (Lee et al., 2007; Woo et al., 2012). To induce Ca^2+^-dependent release from an astrocyte, TFLLR, an agonist of protease-activated receptor-1 (PAR-1), was locally applied, and sensor current elicited by a release of glycine or D-serine was recorded from a HEK293T cell expressing a biosensor (Figure 2A). To discriminate between glycine and D-serine, we utilized either NMDAR (NR1-1a and chimeric NR2A(2D-S1)) (Chen et al., 2008) for the detection of both glutamate and co-agonist, or glycine receptor (human glycine receptor alpha 1; hGlyR α1 L261F) (Laube et al., 2000) for the detection of glycine, but not D-serine, as the biosensor (Figure 2B). When astrocytic Ca^2+^ was elicited by TFLLR, we observed a significant NMDAR-sensor-current, whereas glycine-receptor-sensor-current was minimally observed (Figures 2C and 2D), indicating that an astrocyte releases glutamate together with an NMDAR co-agonist other than glycine, most likely D-serine. To test whether this NMDAR co-agonist is indeed D-serine, we expressed shRNA for serine racemase (SR), D-serine synthesizing enzyme that converts L-serine to D-serine, to gene-silence SR (Figures 2E, S4A-S4E). SR shRNA-expressing astrocyte showed a significantly decreased NMDAR-sensor-current compared to control shRNA-expressing astrocyte (Figure 2E). This decreased NMDAR current was recovered by a 5-minute incubation with 100μM D-serine on the same cell (Figure 2F). These results demonstrate that an astrocyte releases D-serine, not glycine, in a Ca^2+^-dependent manner to activate NMDAR.

**Figure 2.**
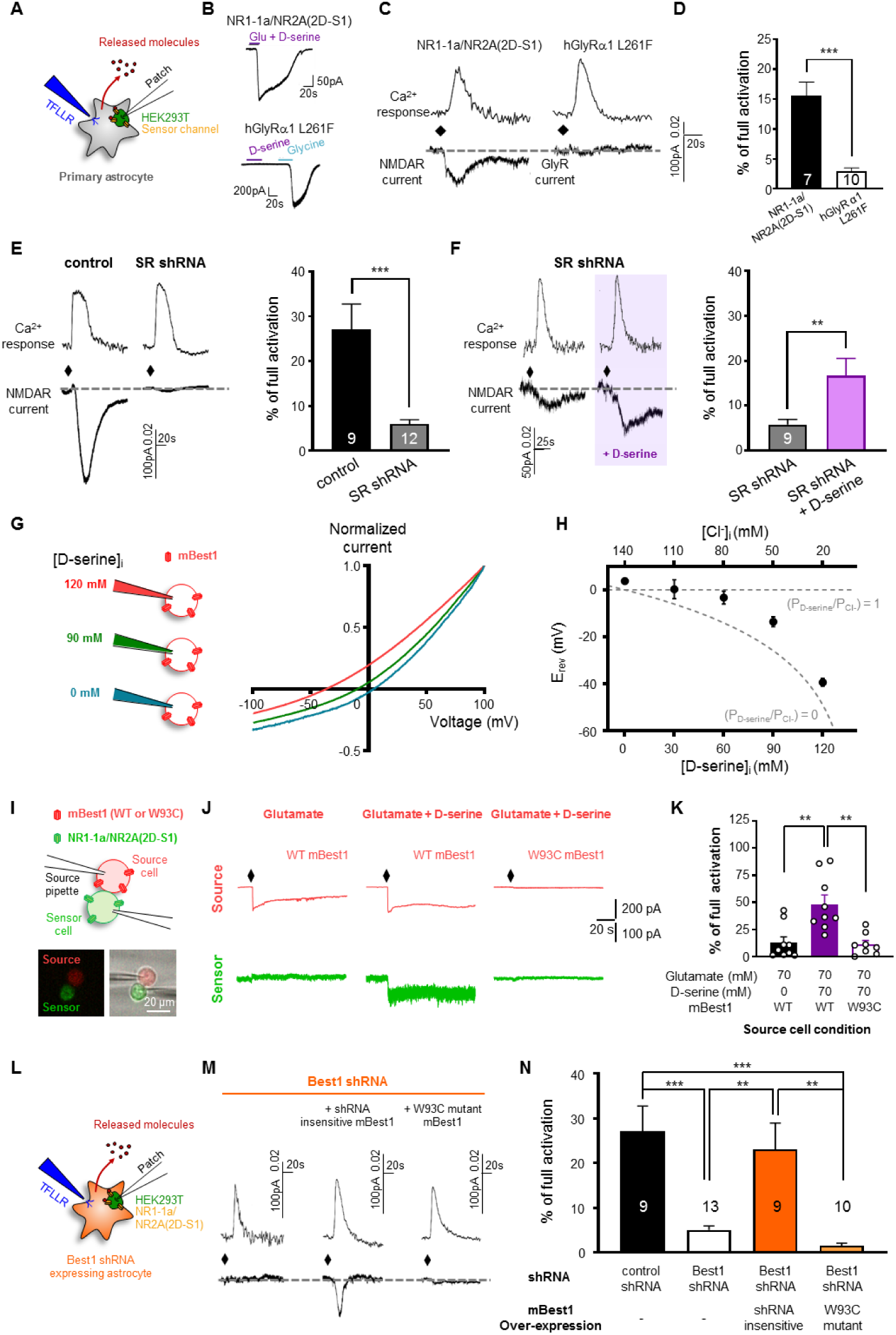
Astrocyte co-releases D-serine and glutamate through Best1 to activate NMDAR. (A) Scheme of sniffer patch using primary astrocyte and sensor cell. (B) Validation of sensor channels. Upper: NR1-1a/NR2A(2D-S1)-mediated responsive current to glutamate and D-serine. Lower: hGlyRα1 L261F-mediated responsive current to D-serine and glycine, respectively. (C) Representative traces of Ca^2+^ response in astrocyte, and responsive sensor current from sensor cell expressing NR1-1a/NR2A(2D-S1) or hGlyRα1 L261F. (D) Summary graph of the peak amplitude normalized to full activation in sensor cell with NR1-1a/NR2A(2D-S1) or hGlyRα1 L261F. (E) Left, representative traces of SR knockdown and control. Right, summary graph of the normalized peak amplitude in each condition. (F) Left, representative traces of SR knockdown before and after 100 µM D-serine treatment. Right, summary graph of the normalized peak amplitude in each condition. (G) I-V relationship in HEK293T cell expressing mouse Best1 (mBest1) in the presence of Ca^2+^ (∼4.5 μM) and varying intracellular concentration of D-serine. (H) Dependence of current reversal potential (Erev, mV) on intracellular D-serine concentration. Gray dotted lines: predicted Erev by the Goldman-Hodgkin-Katz equation when D-serine is as permeable as Cl^−^ (pD-serine/pCl = 1) and when D-serine is not permeable at all (pD-serine/pCl = 0). (I) Scheme of two cell assay. Source cell expressing mBest1 WT or W93C mutant. (J) Representative traces of currents simultaneously recorded from source (red) and sensor (green) cells. (K) Summary graph of the normalized peak amplitude in each condition. (L) Sniffer patch with Best1 shRNA expressing astrocyte. (M) Representative traces of Best1 knockdown without or with either over-expression of shRNA insensitive form or W93C mutant form of Best1. (N) Summary graph of the normalized peak amplitude. Individual dots and numbers refer to cells. Data are represented as mean ± SEM. *p < 0.05; **p < 0.01; ***p < 0.001; unpaired-t test (D and E), paired t-test (F), and one-way ANOVA with Tukey’s multiple comparison test (K and N).

We have previously reported that the astrocytic Ca^2+^-activated anion channel Best1 is permeable to glutamate and mediates the release of glutamate from astrocytes to target synNMDAR (Park et al., 2015; Woo et al., 2012). Thus, we examined a possibility that D-serine could directly permeate Best1. To estimate the relative permeability of D-serine to Best1, we recorded Ca^2+^-activated Best1-mediated current with serial substitutions of chloride in internal pipette solution with equivalent concentrations of D-serine (Figures 2G and 2H), as previously described for glutamate (Park et al., 2009). The gray dotted lines indicate the predicted reversal potential, E_rev_ by the Goldman-Hodgkin-Katz equation when D-serine is as permeable as Cl^−^ (P_D-serine_/P_Cl_ = 1) and when D-serine is not permeable at all (P_D-serine_/p_Cl_ = 0) (Figure 2H). We found that the observed E_rev_’s fell somewhere in between the two dotted lines, indicating that D-serine permeability is greater than zero but less than that of chloride (Figure 2H). At the intracellular D-serine concentration of 90mM, the permeability ratio of D-serine to chloride (P_D-serine_/P_Cl_^-^) was estimated as 0.69, according to the Goldman-Hodgkin-Kats equation (Figure 2H). These results indicate that Best1 has a substantial permeability to D-serine.

Considering the fact that Best1 is permeable to both D-serine and glutamate, we next asked whether D-serine and glutamate can be co-released through Best1. To directly test this possibility, we employed two-cell-sniffer-patch technique, consisting of a source cell (HEK293T cell) expressing the full-clone of mouse *Best1* (*mBest1*) and a sensor cell (HEK293T cell) expressing NR1/NR2A(2D-S1) (Figure 2I). Pre-rupture configuration for the source cell was prepared by forming a gigaseal with an internal pipette solution containing 70mM glutamate with or without 70mM D-serine, and corresponding sensor current was measured during a membrane-rupture of the source cell. For an immediate activation of mBest1 in the source cell upon membrane-rupture, the internal pipette solution contained 4.5 μM free Ca^2+^. We found that upon the membrane rupture of the source cell expressing WT mBest1, a significant sensor NMDAR-current was observed only when the source-cell internal pipette solution contained both glutamate and D-serine (Figure 2J and 2K), indicating that Best1 releases both glutamate and D-serine simultaneously. This Best1-mediated co-release of glutamate and D-serine was absent when the source cell expressed mBest1-W93C, a pore-mutant form of mBest1 (Figure 2J and 2K), further strengthening the concept of a co-release of glutamate and D-serine through Best1.

Finally, we tested the concept of co-release of glutamate and D-serine through Best1 in astrocytes. To test the idea, sniffer-patch experiment was performed with Best1 shRNA-expressing cultured solitary astrocytes (Figures 2L). Best1 shRNA-expressing astrocytes showed almost complete elimination of the sensor-NMDAR-current, which was fully reconstituted by a co-expression of shRNA-insensitive form of mBest1, whereas co-expression of shRNA-insensitive mBest1-W93C showed no recovery (Figure 2M and 2N). Taken together, these results suggest that astrocytes co-release D-serine and glutamate through Best1 in a Ca^2+^-dependent manner to activate adjacent NMDAR and mediate NMDAR tone in the hippocampus.

### Decreased NMDAR tone leads to impaired LTD in hippocampus

Equipped with the molecular and genetic tools to regulate NMDAR tone via Best1, we examined the potential role of NMDAR tone in synaptic plasticity. To assess synaptic plasticity, we performed field EPSP (fEPSP) recordings of the Schaffer collateral pathway at CA3-CA1 synapses, as previously described (Nam et al., 2019; Park et al., 2015). We found that low frequency stimulation (LFS, 900 stimuli at 1 Hz)-induced LTD was completely impaired in Best1 KO (Figures 3A-3C, S5C and S5D), whereas effects produced by high frequency stimulation (HFS, 100 stimuli at 100 Hz) or 10 Hz stimulation (900 stimuli at 10 Hz) showed no difference between WT and Best1 KO (Figures 3C, S5A and S5B). These results suggest that regulation of NMDAR tone through Best1 is critical for the induction of LTD, but not LTP in the hippocampus. To test if the impaired LTD in Best1 KO is due to the lack of Best1-mediated co-release of D-serine and glutamate in astrocytes, we over-expressed Best1 in hippocampal astrocytes by injection of AAV-GFAP-Best1-IRES-EGFP virus into hippocampal CA1 of Best1 KO (Figures 1H-1K, S1D and S1E) and examined LFS-induced LTD (Figures 3D-3F). We found that hippocampal LTD in Best1 KO was fully restored by astrocytic Best1 rescue (GFAP-Best1), but not by control virus (GFAP-GFP) (Figures 3E and 3F). These results indicate that astrocytic Best1 is sufficient for hippocampal LTD, possibly via regulation of NMDAR tone.

**Figure 3.**
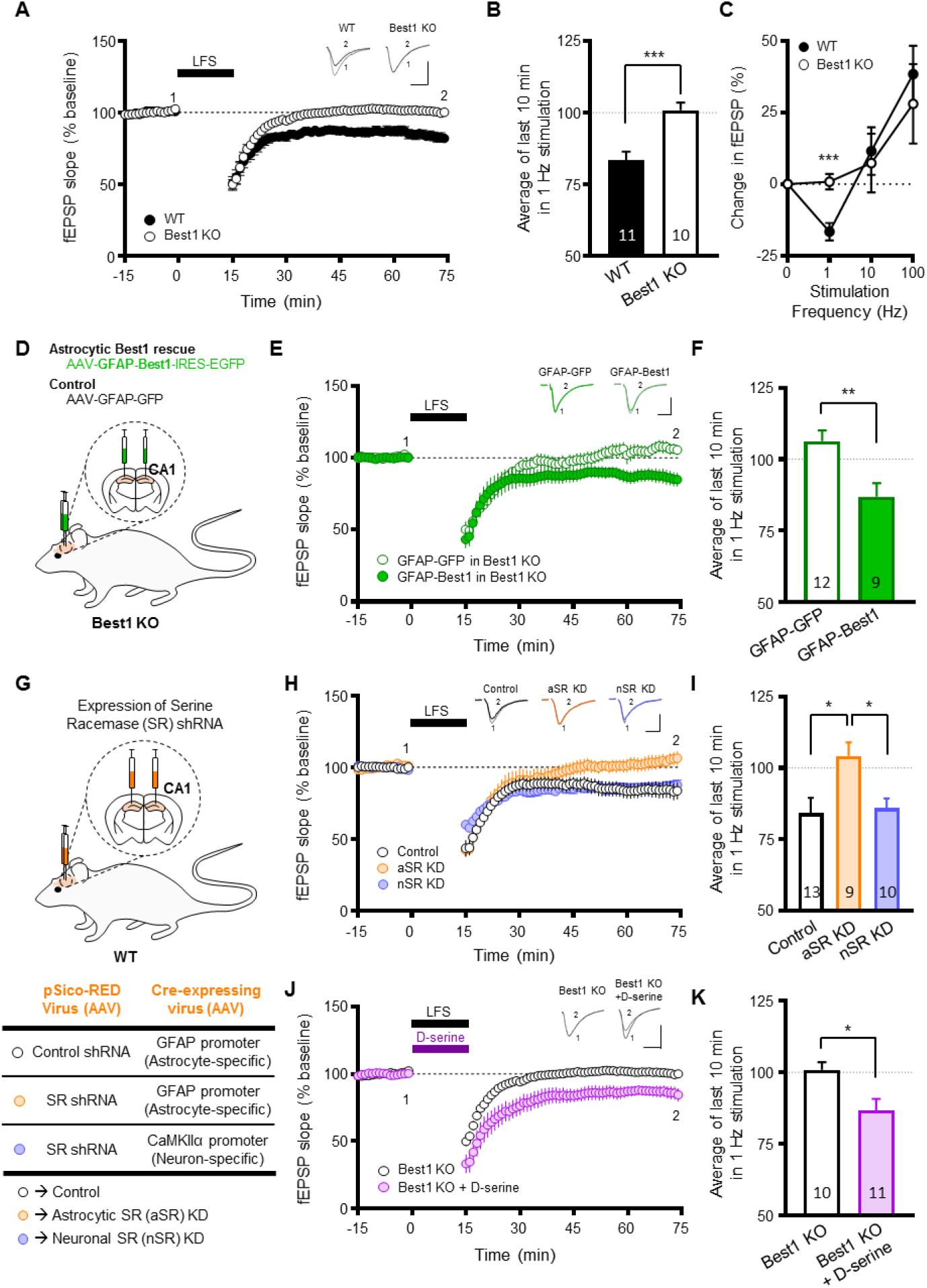
Astrocytic regulation of NMDAR tone through Best1 is important for the induction of LTD. (A) LFS (900 stim. at 1 Hz)-induced LTD in WT and Best1 KO. (B) Summary graph of LFS-induced LTD in each condition. (C) Bienenstock–Cooper–Munro (BCM) curve of synaptic plasticity in each condition. (D) Scheme of CA1 astrocyte-specific Best1 rescue in Best1 KO. (E and F) LFS-induced LTD (E) and summary graph (F) in CA1 astrocyte-specific Best1 rescue from Best1 KO. (G) Scheme of cell-type specific SR knockdown. (H and I) LFS-induced LTD (H) and summary graph (I) in cell-type specific mSR knockdown experiment. (J and K) LTD induction in Best1 KO with D-serine application during LFS (J) and summary graph (K). Numbers in the graphs refer to hippocampal slices. Data are represented as mean ± SEM. *p < 0.05; **p < 0.01; ***p < 0.001; unpaired t-test (B, C, F and K), and one-way ANOVA with Tukey’s multiple comparison test (I).

It has been previously reported that depletion of D-serine impairs (Zhang et al., 2008) or reduces LTD (Papouin et al., 2012). However the cellular source of D-serine was not determined. Potential sources of D-serine includes astrocyte (Papouin et al., 2017) and neuron (Wolosker et al., 2017). To examine the cellular source of D-serine required for LTD induction, we inhibited D-serine synthesis in a cell-type specific manner with a Cre recombinase (Cre)-dependent SR shRNA expressing (AAV-pSico-RED SR shRNA) virus, in combination with either astrocyte-specific (AAV-GFAP-Cre) or excitatory neuron-specific (AAV-CaMKIIα-Cre) virus (Figure 3G). We found that astrocytic SR knockdown (aSR KD) eliminated LTD, whereas neuronal SR knockdown (nSR KD) did not (Figures 3H and 3I). These results indicate that astrocytic D-serine, but not neuronal D-serine, is critical for LTD induction. Finally, we tested whether the impaired LTD in Best1 KO can be restored by increasing NMDAR tone during LTD induction. To increase NMDAR tone in Best1 KO, 20 μM D-serine was applied during LFS (Figure 3J). We found that application of D-serine significantly restored the impaired LTD in Best1 KO (Figures 3J and 3K), indicating that NMDAR tone during LFS is critical for hippocampal LTD. Taken together, these results indicate that NMDAR tone mediated by astrocytic D-serine during LFS causes hippocampal LTD.

### NE induces NMDAR tone and LTD in hippocampus

It has been reported that exogenously applied NE by itself (without LFS) induces NMDAR-dependent LTD (NE-LTD) through α1-AR (Scheiderer et al., 2004), which is predominantly expressed in astrocytes (Hertz et al., 2010). To test if NE-LTD is caused by NMDAR tone, we first examined whether NE can increase astrocytic Ca^2+^ through α1-AR and NMDAR tone via Best1 in the hippocampus. We expressed jRCaMP1a, a genetically encoded Ca^2+^ indicator, in CA1 astrocytes with AAV-GFAP104-jRCaMP1a virus (Figures 4A and 4B) and observed robust Ca^2+^ responses in astrocytes by 200 μM NE application, which were significantly blocked by prazosin, an α1-AR blocker (Figure 4C). These results indicate that NE induces astrocytic Ca^2+^ through α1-AR, consistent with a previous report (Duffy and MacVicar, 1995). We then recorded NE-induced exNMDAR current and found that NE caused a significant exNMDAR current in CA1 pyramidal neuron, which was completely blocked by APV (Figures 4D and 4F). This NE-induced exNMDAR current was almost completely eliminated in Best1 KO, and significantly restored by D-serine application (Figures 4E and 4F), indicating that NE increases NMDAR tone through Best1 in the hippocampus. To investigate whether NE-LTD is mediated by NMDAR tone through Best1, we performed fEPSP recordings in WT and Best1 KO, and found that NE-induced LTD was blocked by APV in WT (Figures 4G and 4I), absent in Best1 KO (Figures 4H and 4I), and restored by D-serine in Best1 KO (Figures 4H and 4I). Taken together, these results indicate that NE activates astrocytic α1-AR to induce LTD by increasing NMDAR tone through Best1.

**Figure 4.**
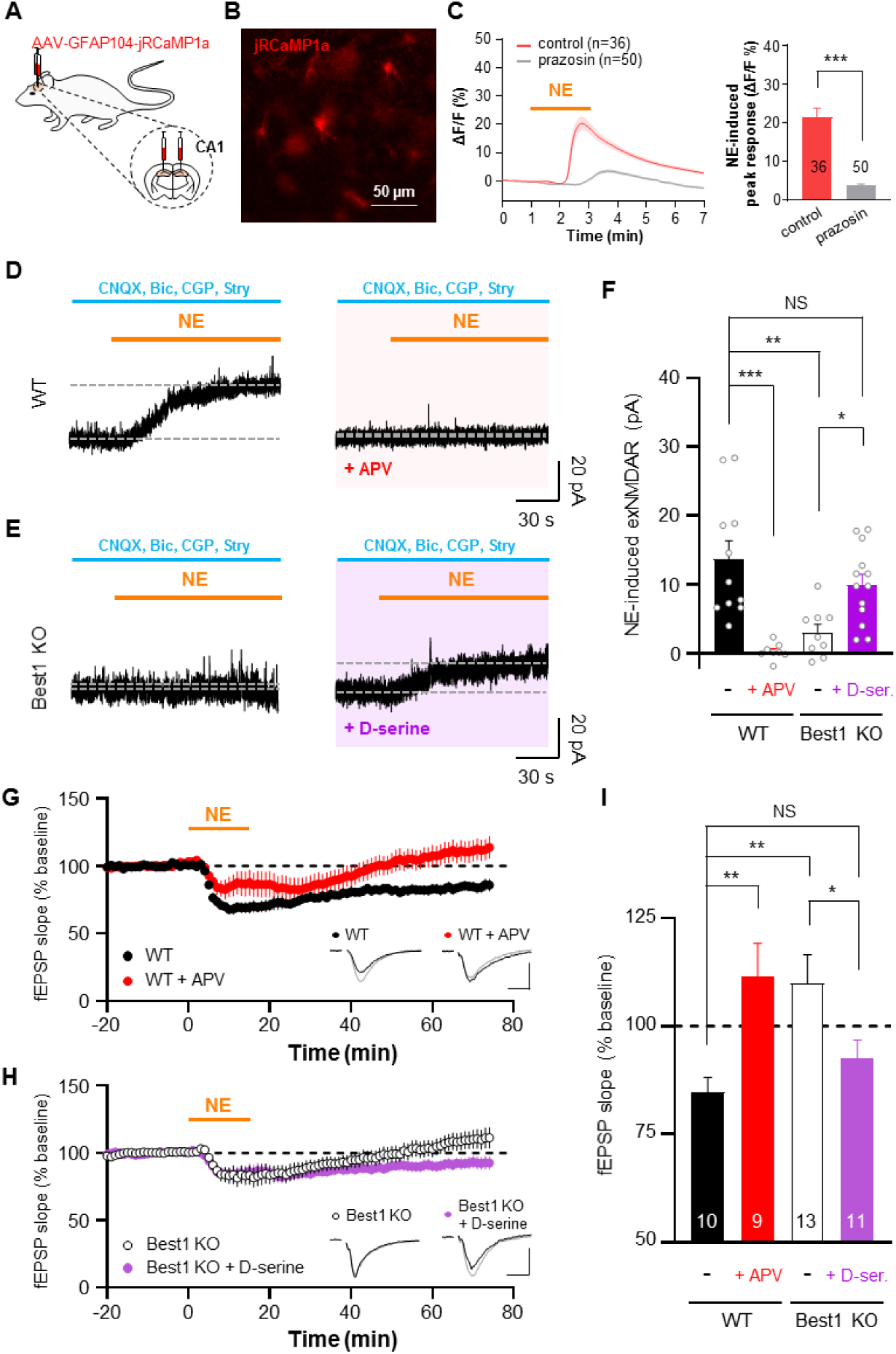
NE induces astrocytic Ca^2+^, NMDAR tone increase and NE-LTD. Astrocytic expression of jRCaMP1a with AAV-GFAP104-jRCaMP1a. (B) Fluorescent image of jRCaMP1a in hippocampal CA1. (C) Left, 200 µM NE-induced jRCaMP1a response with or without 10 µM prazosin, α1-AR blocker. Right, Summary of NE-induced peak response. (D) Representative traces of NE-induced exNMDAR current in WT with or without APV. (E) Representative traces of NE-induced exNMDAR current in Best1 KO with or without 100 µM D-serine. (F) Summary graph of NE-induced exNMDAR current in each condition. (G) NE-induced LTD in WT with or without APV. (H) NE-induced LTD in Best1 KO with or without 100 µM D-serine. (I) Summary graph of NE-induced LTD in each condition. Numbers in the graphs refer to cells (in C) or hippocampal slices (in I), and individual dots refer to cells. Data are represented as mean ± SEM. *p < 0.05; **p < 0.01; ***p < 0.001; Mann Whitney test (C), One-way ANOVA with Tukey’s multiple comparisons test (F), and unpaired t-test (I).

### Endogenous NE is released via axo-axonic synapses to mediate heterosynaptic LTD

The release of NE throughout the brain is critical for modulating arousal, attention, and cognitive behaviors, and its disruption is strongly associated with several psychiatric and neurodegenerative disorders in humans (Schwarz and Luo, 2015). Most of the NE released in the brain is supplied by the fiber projections from a very small, bilateral nucleus in the brainstem called the locus coeruleus. The local release of NE is reported to be stimulated by glutamate (Howells and Russell, 2008; Malva et al., 1994) or conventional electrical stimulation of Schaffer collateral pathway (Feng et al., 2019), raising a possibility that local NE release (Jacobowitz, 1979) is mediated by axo-axonic synapses in the hippocampus (Schwarz and Luo, 2015). Thus, we investigated the local NE release onto hippocampal astrocytes and its role for LTD. To visualize local NE release onto hippocampal astrocytes, GRAB_NE2m_, a GPCR-based NE fluorescence sensor, was expressed in hippocampal astrocytes via injection of AAV-GFAP104-GRAB_NE2m_ into CA1 (Figures 5A and 5B), and Schaffer collateral pathway was stimulated with various stimulation intensities and frequencies (Figures 5C and 5D). We found that the electrical stimulation increased GRAB_NE2m_ fluorescence from astrocytes, in stimulation intensity- and frequency-dependent manners, with a peak response at 50 Hz (Figures 5E-5G). The evoked fluorescence was significantly blocked by Yohimbine (Figure 5H), indicating that NE is released onto hippocampal astrocytes during the stimulation of Schaffer collateral pathway. To test whether glutamate contributes to the local NE release onto astrocytes, blockers of glutamate receptors were independently applied during the stimulation. We found that the NE response was majorly blocked by inhibition of AMPAR with CNQX, but minimally by inhibition of mGluR5 with MPEP or NMDAR with APV (Figures 5I-5K). These results indicate that stimulation of Schaffer collateral pathway induces local NE release by activating AMPAR, most likely localized at the axo-axonic presynaptic terminals of the LC projection fibers (Schwarz and Luo, 2015).

**Figure 5.**
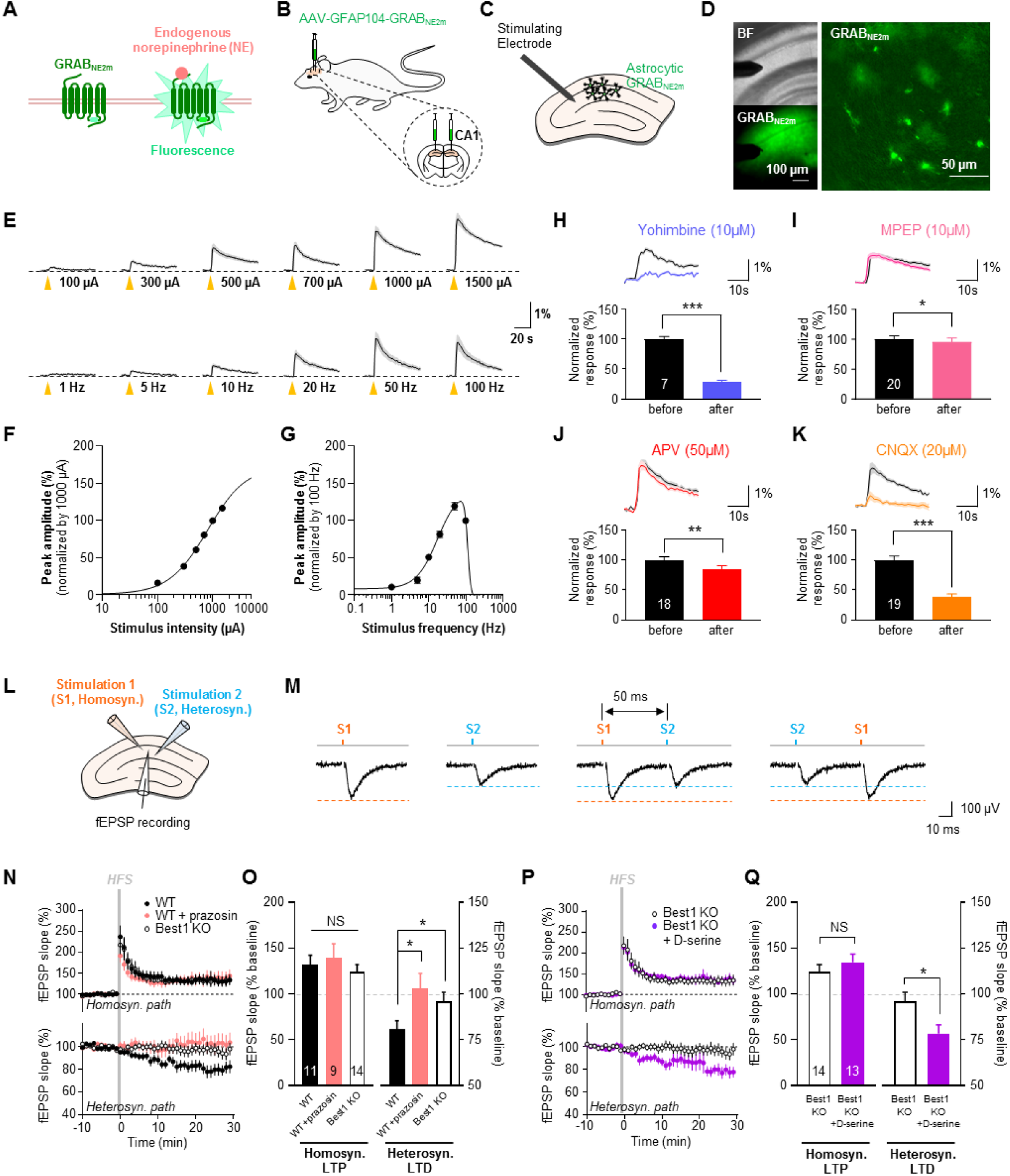
Endogenously released NE by glutamate induces heterosynaptic LTD through astrocytes. (A) GRABNE2m, a fluorescent sensor for NE. (B) Astrocytic expression of GRABNE2m with AAV-GFAP104-GRABNE2m. (C) Scheme of evoked NE release by Schaffer collaterals stimulation. (D) Images of GRABNE2m in hippocampal CA1. (E) Representative traces of GRABNE2m response by the various stimulation. (F) Stimulus intensity-GRABNE2m response curve (at 20 Hz). (G) Stimulus frequency-GRABNE2m response curve (at 500 μA). (H-K) Inhibition of GRABNE2m response by Yohimbine (H), MPEP (I), APV (J) and CNQX (K). (L) Scheme of simultaneous homosynaptic (S1, orange) and heterosynaptic (S2, blue) recordings. (M) Lack of heterosynaptic facilitation with 50 ms interval. (N) Homosynaptic and heterosynaptic changes by HFS in WT, WT with 10 µM prazosin, and Best1 KO. (O) Summary graph of fEPSP changes in (N). (P) Homosynaptic and heterosynaptic changes by HFS in Best1 KO and Best1 KO with 100 µM D-serine. (Q) Summary graph of fEPSP changes in (P). Numbers in the graphs refer to hippocampal slices. Data are represented as mean ± SEM. *p < 0.05; **p < 0.01; ***p < 0.001; unpaired t-test (O and Q) and paired t-test (H, I, J, K).

It is worth noting that the local NE release was prominent at HFS (Figures 5E-5G), which causes not only homosynaptic LTP but also heterosynaptic LTD (Scanziani et al., 1996). To investigate the role of local NE release in homosynaptic LTP and heterosynaptic LTD, we performed simultaneous recordings of homosynaptic and heterosynaptic fEPSP (Figure 5L). Two independent pathways were accessed with theta micropipette as a bipolar microelectrode and validated by the lack of heterosynaptic paired-pulse facilitation (Figure 5M). We observed a robust induction of heterosynaptic LTD while homosynaptic LTP was induced by HFS (Figures 5N and 5O). Surprisingly, the heterosynaptic LTD, but not homosynaptic LTP, was blocked by prazosin (Figures 5N and 5O), indicating that α1-AR activation is necessary for heterosynaptic LTD. More importantly, we observed that the heterosynaptic LTD, but not homosynaptic LTP, was impaired in Best1 KO (Figures 5P and 5Q). The impaired heterosynaptic LTD in Best1 KO was fully restored by an enhancement of NMDAR tone with D-serine (Figures 5P and 5Q). Taken together, these results indicate that local NE-α1-AR signaling mediates heterosynaptic LTD through NMDAR tone.

### NMDAR tone-dependent heterosynaptic LTD is required for repotentiation LTP

Heterosynaptic plasticity has been proposed to enable further changes in synaptic plasticity, i.e., metaplasticity (Chen et al., 2013b). Given that Best1 KO showed the lack of heterosynaptic LTD, we examined further changes in synaptic plasticity after HFS in Best1 KO. To test a bidirectional modification of metaplasticity after the 1^st^ HFS potentiation LTP, we delivered additional LFS and 2^nd^ HFS during fEPSP recordings (Figure 6A), as previously described (Dudek and Bear, 1993). We found that the LFS-induced depotentiation LTD was observed in both WT and Best1 KO (Figures 6A and 6B). In contrast, the 2^nd^ HFS-induced repotentiation LTP was significantly impaired in Best1 KO (Figures 6A and 6B). This pattern of intact depotentiation LTD and impaired repotentiation LTP in Best1 KO was similarly observed when LTP-inducing stimulation was changed from HFS to theta-burst stimulation (TBS) (Figures S5E and S5F). These results indicate that the lack of NMDAR tone and heterosynaptic LTD in Best1 KO leads to the impaired repotentiation LTP. To test whether the impaired repotentiation LTP in Best1 KO can be restored by an enhancement of NMDAR tone during each stimulation period, we applied D-serine during 1^st^ HFS potentiation (orange), LFS (green), or 2^nd^ HFS repotentiation (blue) (Figures 6C and 6E). We found that the impaired repotentiation LTP in Best1 KO was fully restored by D-serine treatment only during 1^st^ HFS potentiation (Figures 6C and 6D), but not during LFS or 2^nd^ HFS repotentiation (Figures 6E and 6F). These results indicate that NMDAR tone-dependent heterosynaptic LTD is critical for subsequent repotentiation LTP and metaplasticity.

**Figure 6.**
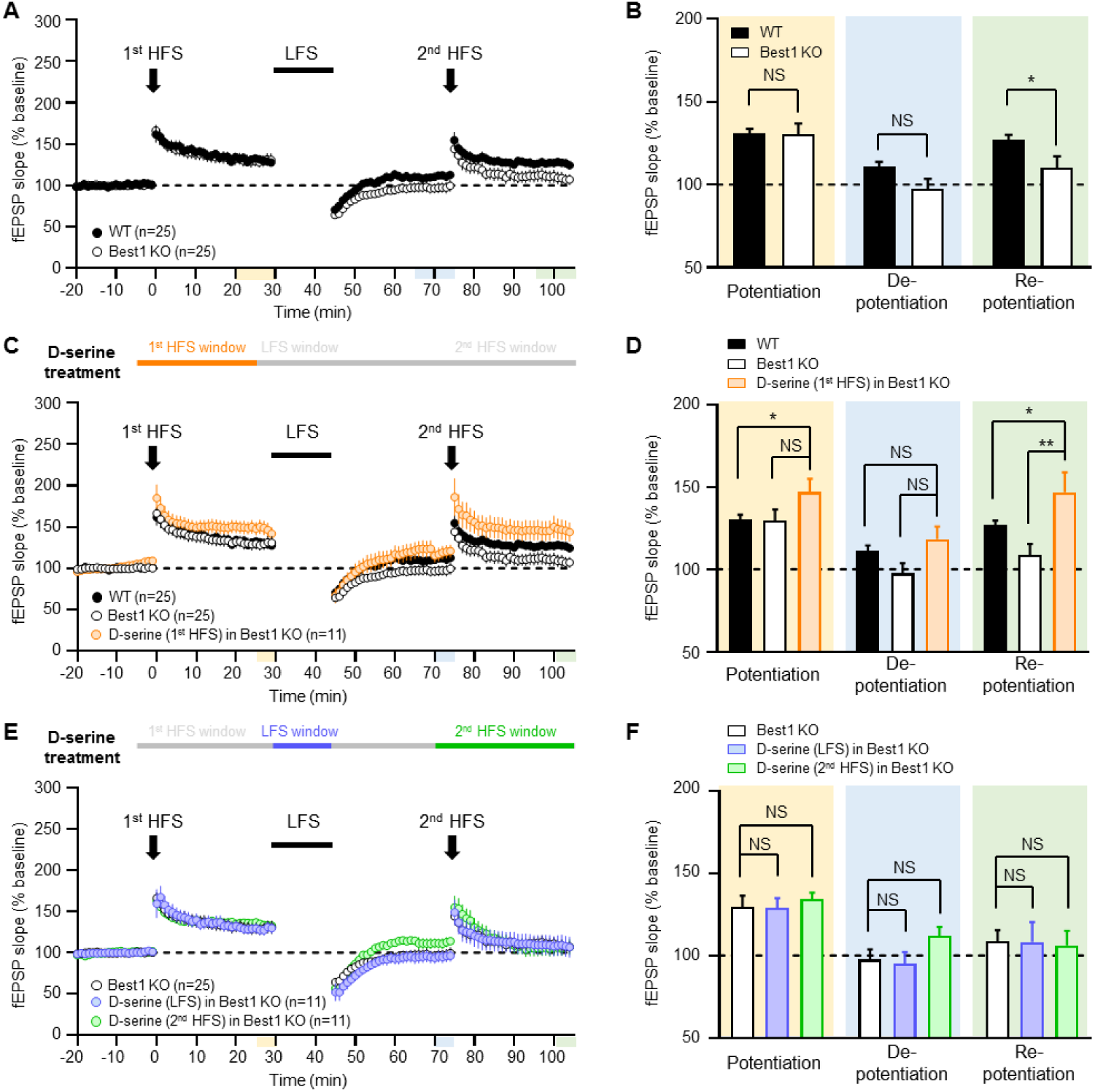
NMDAR tone during first potentiation facilitate repotentiation. (A and B) Time course of the normalized fEPSP slope changes (A) and summary graph (B) of 1^st^ HFS-induced potentiation, LFS-induced depotentiation and 2^nd^ HFS-induced repotentiation in WT and Best1 KO. (C) NMDAR tone enhancement during 1^st^ HFS window in Best1 KO by 20 µM D-serine treatment. (D) Summary graph of results from (C). (E) NMDAR tone enhancement during LFS or 2^nd^ HFS window in Best1 KO by 20 µM D-serine treatment. (F) Summary graph of the results from (E). Data are represented as mean ± SEM. *p < 0.05; **p < 0.01; ***p < 0.001; unpaired t-test (B, D and F).

### Spatial reversal learning is impaired in Best1 KO and rescued by astrocytic Best1

To investigate the role of NMDAR tone-dependent heterosynaptic LTD in learning and memory, we performed various hippocampus-dependent memory tasks such as Morris water maze (MWM) test, passive avoidance test (PAT) and contextual fear conditioning (CFC) test with WT and Best1 KO (Figures 7A and S6B-S6E). We found that in MWM, PAT, and CFC, there was no difference in memory acquisition between WT and Best1 KO (Figure 7B and Figures S6B-S6E). In contrast, we observed that Best1 KO showed significantly impaired spatial reversal learning when the hidden platform was relocated at opposite quadrant (O) during the reversal task session (Figures 7B and 7C); Best1 KO spent significantly more time in the original target quadrant (O) and significantly less time in T, compared to WT (Figure 7C). This impaired spatial reversal learning in Best1 KO was not due to a malfunction of vision or locomotion, as evidenced by an intact acquisition when we switched from the hidden to a visible platform (Figures 7D, 7E and S6A). These results imply that Best1-dependent NMDAR tone and heterosynaptic LTD are necessary for spatial reversal learning and flexible memory. To further test whether astrocyte-specific Best1 rescue sufficiently restores spatial reversal learning in Best1 KO, we injected AAV-GFAP-Best1-IRES-EGFP virus bilaterally into CA1 hippocampus of Best1 KO and performed MWM test. We found a significant restoration of spatial reversal learning in Best1 KO (Figure 7F), indicating that CA1-astrocyte-specific Best1 rescue is sufficient for spatial reversal learning and flexible memory. Taken together, these results establish a causal relationship between the astrocytic Best1 in CA1 hippocampus and spatial reversal learning and flexible memory.

**Figure 7.**
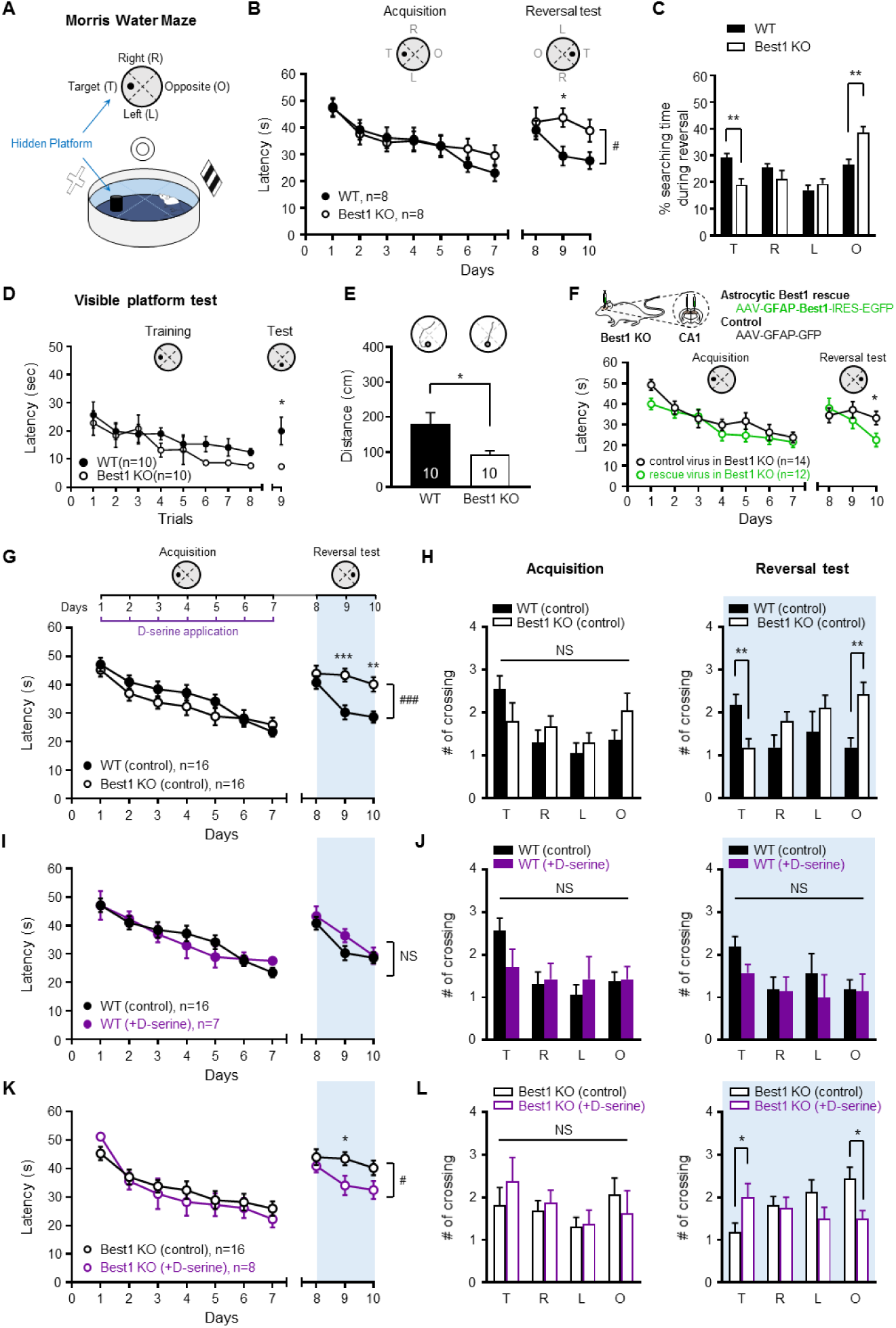
NMDAR tone is critical for formation of flexible memory. (A) Scheme of Morris water maze. (B) Escape latency of WT and Best1 KO in acquisition and reversal test session. (C) Searching time of WT and Best1 KO during reversal test in each quadrant. (D and E) Escape latency (D) and moved distance (E) of WT and Best1 KO in visible platform test. (F) Escape latency of Best1 KO with the rescue of astrocytic Best1, in acquisition and reversal test session during hidden platform test. (G-L) Application of saline or D-serine (600 mg/kg) in each day of acquisition session. (G), (I), (K) Escape latency of WT and Best1 KO with application of saline or D-serine. (H), (J), (L) Left: number of crossing to each quadrant in each condition during acquisition session. Right: number of crossing to each quadrant in each condition during reversal test session. Numbers in the graphs refer to animals. Data are represented as mean ± SEM. *p < 0.05; **p < 0.01; ***p < 0.001; Two-way ANOVA with Fisher’s LSD (B, D, F) and unpaired t-test (C, E, G-L). #p < 0.05; ##p < 0.01; ###p < 0.001; Two-way repeated measures (RM) ANOVA (genotype or treatment) (B, G, I, K).

### NMDAR tone increase during memory acquisition is critical for flexible memory

Although LFS-induced homosynaptic LTD has been implicated in spatial reversal learning (Dong et al., 2013; Nicholls et al., 2008), there have been conflicting results (Rutten et al., 2011; Zhang and Wang, 2013), raising a possibility that heterosynaptic, not homosynaptic, LTD might be responsible for spatial reversal learning. Considering the fact that heterosynaptic LTD occurs at the time of homosynaptic LTP, we hypothesized that heterosynaptic LTD during the initial memory acquisition contributes to spatial reversal learning and flexible memory, as it enables further repotentiation LTP (Figures 6C and 6D). To test this hypothesis, Best1 KO mice were injected with D-serine (600 mg/kg, intra-peritoneal) during the initial memory acquisition session in MWM (Figures 7G-7L). We found that application of D-serine during the initial memory acquisition session fully restored the impaired spatial reversal learning in Best1 KO (Figures 7K and 7L). These results imply that heterosynaptic LTD during the initial memory acquisition is critical for the formation of flexible memory (Figures S7A-S7D). Altogether, these findings establish that astrocytes render memory flexible through regulation of Best1-dependent NMDAR tone and heterosynaptic LTD during the initial stage of memory acquisition.

## Discussion

In the present study, we have demonstrated for the first time that astrocytes are critically involved in reversal learning and flexible memory. Astrocytes achieve this unique function by co-releasing D-serine and glutamate through the Ca^2+^-activated anion channel Best1 upon activation of NE-α1-AR signaling pathway, leading to an enhanced NMDAR tone and induction of heterosynaptic LTD during the initial period of memory acquisition (Figure S7A). The presence of astrocyte-driven NMDAR tone provides the molecular basis for both the formation of a flexible memory at the time of initial memory acquisition and the induction of repotentiation LTP and a new memory at the time of reversal learning (Figure S7B). In the absence of such astrocyte-derived NMDAR tone as in Best1 KO mice, the animals exhibited an impaired heterosynaptic LTD, impaired metaplasticity (*i.e.*, repotentiation LTP), impaired reversal learning and flexible memory (*i.e.*, a persistence of initial memory) (Figure S7C), which were all restored by a D-serine supplement or the astrocyte-specific reconstitution of Best1 in Best1 KO mice (Figure S7D). The impaired reversal learning and flexible memory are reminiscent of the major symptoms of ASD and schizophrenia, both of which share the common mechanism of NMDAR hypofunction. The unexpected role of NE and the astrocytic α1-AR in the induction of heterosynaptic LTD is not surprising, considering the fact that NE is critical for attention, focus, and arousal. Our research will provide a comprehensive understanding of NE, NMDAR tone, and memory formation.

### Astrocyte as a regulator of NMDAR tone

Since the first discovery of tonic activation of exNMDAR in CA1 pyramidal neurons (Sah et al., 1989), many independent research groups have investigated the molecular and cellular sources and functions of the tonic activation of exNMDAR using various nomenclatures such as “tonic NMDAR” and “NMDAR tone” (Parsons and Raymond, 2014). Although the two names may sound indistinguishable, they have profoundly different implications if one considers the properties of NMDAR. For a channel activation of exNMDAR, glutamate and co-agonist (glycine or D-serine) have to bind to their respective binding site at exNMDAR (Traynelis et al., 2010). However, the binding of two agonists is not sufficient for tonic activation of exNMDAR because of the Mg^2+^ block near resting membrane potential. The glutamate-and-co-agonist-bound exNMDAR is activated only when a depolarization relieves the Mg^2+^ block (Sah et al., ^1^98^9^). This is in great contrast to the extrasynaptic GABA_A_ receptors (GABA_A_R), in which the receptors are tonically activated voltage-independently upon the binding of GABA (Egawa and Fukuda, 2013). Thus, it is appropriate to name it, “tonic GABA_A_R.” In contrast, for the exNMDAR in which glutamate and co-agonist are bound but not active under resting membrane potential, it is more appropriate to use the term “NMDAR tone”, rather than “tonic NMDAR”, as we did in the present study.

At first, astrocytes have been proposed to mediate NMDAR tone in the hippocampus (Le Meur et al., 2007). However, this concept of astrocytic contribution to NMDAR tone has been challenged by the report that knockout of *IP3R2* (type 2 inositol 1,4,5-trisphosphate receptor), which is known to mediate endoplasmic reticulum (ER) Ca^2+^ release exclusively in astrocytes (Holtzclaw et al., 2002), showed no difference in hippocampal NMDAR tone (Petravicz et al., 2008). Contrary to this conflicting observation, we have found that astrocytes majorly contribute to the hippocampal NMDAR tone in a Ca^2+^-dependent manner through the Ca^2+^-activated anion channel Best1 (Figure 1). Our results raise a possibility that IP3R2-mediated Ca^2+^ signal may not be necessary for the activation of Best1. In support of this possibility, IP3R2-independent Ca^2+^ release from ER has been reported (Okubo et al., 2018), and, more importantly, ER-Ca^2+^-independent Ca^2+^ signal is found in astrocytic fine processes (Rungta et al., 2016; Srinivasan et al., 2015), where Best1 is mostly expressed (Woo et al., 2012). Other Ca^2+^ sources (e.g., spotty Ca^2+^ from TRPA1) (Oh et al., 2019; Shigetomi et al., 2013) could also contribute to the activation of Best1, and should be investigated in the future studies.

In a series of previous studies, we have demonstrated that 1) Best1 is permeable to glutamate and mediates astrocytic glutamate release upon activation of PAR1 (Oh et al., 2012; Park et al., 2015; Park et al., 2013; Park et al., 2009; Woo et al., 2012), 2) PAR1-driven astrocytic glutamate release targets postsynaptic NMDARs, and 3) modulates hippocampal synaptic plasticity by lowering the threshold for LTP (Lee et al., 2007; Park et al., 2015). In the current study, we have further identified D-serine as a novel permeant anion passing through Best1 (Figure 2), and demonstrated that astrocytic Best1 is an ideal regulator of NMDAR tone by releasing both glutamate and D-serine in hippocampus. Our study directly addresses the recent controversy over the origin of D-serine (Papouin et al., 2017; Wolosker et al., 2017) and provides answers by the astrocyte-specific manipulation of D-serine release through over-expression of Best1 in astrocytes of the Best1 KO mice. Our results are consistent with the previous reports that 1) D-serine affects not only LTP, but also LTD induction (Zhang et al., 2008), and 2) inhibition of the mechanism mediating neuronal D-serine release does not affect LTD (Sason et al., 2017). Additionally, utilizing the cell-type specific expression of SR shRNA, we have demonstrated that D-serine synthesis from astrocytes, but not neurons, is necessary for LTD induction (Figure 3). These results indicate that both co-release of D-serine and glutamate from astrocyte through Best1 and D-serine synthesis by astrocytic SR are critical for hippocampal LTD.

Interestingly, we have found that D-serine administration alone was able to restore the impairments of Best1 KO. These results appear to underrate the role of Best1-mediated glutamate release in the hippocampus. However, under physiological condition Best1-mediated glutamate also contributes to NMDAR tone (Figure 1), and is expected to play an important role when PAR1 is activated (Park et al., 2015; Price et al., 2021). Indeed, we could observe that ambient glutamate was decreased by 37.4% in Best1 KO compared to WT (Figure 1M), implying that there is a substantial remaining portion of ambient glutamate. These results suggest that, in addition to Best1-mediated tonic glutamate release, there may exist other alternative mechanisms for tonic glutamate release. Swell1 (Yang et al., 2019) and Ttyh (Han et al., 2019), recently identified as astrocytic VRAC (volume-regulated anion channel), can be alternative mechanisms for glutamate release. In addition, astrocytic vesicular glutamate release (Araque et al., 2000), or cystine-glutamate antiporter xCT (SLC7A11) (Ottestad-Hansen et al., 2018) may also contribute to NMDAR tone. These exciting possibilities await future investigations.

As astrocytes are heterogeneous in different brain regions (Khakh and Sofroniew, 2015), how the regulatory mechanism of NMDAR tone or intracellular metabolites differ from one brain region to another should be further investigated to reveal differential roles of NMDAR tone in different brain regions. It should be noted that Best1 mediates glutamate and D-serine release in the cortex as well **(Lalo et al., 2021)**, suggesting that it may also mediate NMDAR tone in medial prefrontal cortex (mPFC) (Povysheva and Johnson, 2012) or other brain regions. Consequently, future researches on different regulatory mechanisms in various brain regions will greatly broaden our understanding of the critical roles of NMDAR tone.

### Local NE release mediates heterosynaptic LTD via astrocytic regulation of NMDAR tone

Activation of LC has been implicated in arousal, attention, and cognitive behaviors (Schwarz and Luo, 2015). However, since activation of LC exerts effects on a large area throughout the brain (Zerbi et al., 2019), an alternative mechanism is required when NE is utilized in local area. In this study, we have found that the glutamate from Schaffer collateral fibers stimulates NE release by activating presynaptic AMPAR (Ghersi et al., 2003) at the LC terminals of so-called en passant varicosities (Atzori et al., 2016) as in the form of axo-axonic synapses. These results support the previously proposed hypothesis for the local control of NE release (Jacobowitz, 1979), and are consistent with the results that glutamate stimulates local NE release (Howells and Russell, 2008; Malva et al., 1994). This local NE release can be the basis of the recently proposed “glutamate amplifies noradrenergic effects” (GANE) model (Mather et al., 2016). Considering the previous report that NE-induced NMDAR-dependent LTD is independent of LFS-induced LTD (Scheiderer et al., 2004), the finding that NE release is prominent only at high frequency, rather than low frequency, stimulation of the Schaffer collateral fibers from the CA3 glutamatergic neurons is not unexpected. High-frequency stimulation of Schaffer collateral fibers induces homosynaptic LTP at stimulated synapses, and simultaneously releases NE to turn-on the cascade of events of 1) activating astrocytic α1-AR, 2) increase in NMDAR tone, and 3) induction of heterosynaptic LTD at unstimulated synapses (Figures 4 and 5). These findings are consistent with the previous reports that astrocytes play an essential role in heterosynaptic LTD (Chen et al., 2013a; Serrano et al., 2006). Our study demonstrates that activation of astrocytes by local NE release subsequently enables the plasticity of neighboring synapses (Figure 6), which goes in line with the concept of astrocytes orchestrating synaptic dynamics (De Pitta et al., 2016). Given the fact that one astrocyte is in contact with about 140,000 synapses from numerous neurons in CA1 of the adult rat (Bushong et al., 2002), it is plausible to consider one astrocyte to mediate heterosynaptic LTD at the unstimulated synapses while the stimulated synapses are potentiated. Thus, astrocytes provide a unique structural medium for a simultaneous dynamic control of multiple synapses, from both stimulated and unstimulated neurons, mediating various forms of homeostatic plasticity and metaplasticity. This notion is also supported by the results of accompanying paper **(Lalo et al., 2021)**.

### Heterosynaptic LTD determines flexibility of memory

Our study attempts to fill in the huge gap between the inadequately simple concept of homosynaptic plasticity and the complex nature of memory formation, retention, and flexibility. Cognitive flexibility has long been explained only by homosynaptic LTD (Dong et al., 2013; Kim et al., 2011; Nicholls et al., 2008). However, because several studies have shown that homosynaptic LTD is not indispensable for cognitive flexibility (Rutten et al., 2011; Zhang and Wang, 2013), an alternative mechanism of synaptic plasticity for cognitive flexibility has been needed. In the present study, we have demonstrated for the first time that heterosynaptic LTD accompanying homosynaptic LTP contributes to cognitive flexibility. The biggest difference in the function between homosynaptic and heterosynaptic LTD for cognitive flexibility is that homosynaptic LTD acts when memory modification is required (Dong et al., 2013), whereas heterosynaptic LTD occurs during memory acquisition. This novel concept is supported by the observation that impaired reversal learning was restored by increasing the NMDAR tone of Best1 KO mice during the initial memory formation (Figure 7). We interpret these results as when the initial memory is formed, the memory that accompanies heterosynaptic LTD becomes “flexible memory”, and the memory that does not accompany becomes “inflexible memory”. This novel idea predicts that less-flexible memory can be formed during memory acquisition under certain conditions in which heterosynaptic LTD is impaired. In support of this prediction, prazosin administered during threat memory formation in mice and humans has been reported to interfere with subsequent extinction learning (Do-Monte et al., 2010; Homan et al., 2017), which is consistent with our observation that impaired heterosynaptic LTD interferes subsequent learning (Figure 7). The decrease in glutamate-induced NE release during aging (Dezfuli et al., 2019) also suggests that declined cognitive flexibility in the elderly (Boone et al., 1993) can also be due to impaired NE-α1-AR-dependent heterosynaptic LTD. Similarly, as LC is one of the most vulnerable regions in the progression of AD (Matchett et al., 2021), declined cognitive flexibility in the early stages of AD (Guarino et al., 2018) can be resulted from the impaired heterosynaptic LTD. In addition to AD, although degeneration of LC in ASD and schizophrenia was found to be minimal (Craven et al., 2005; Martchek et al., 2006), the impaired cognitive flexibility is possibly due to a decreased local NE release or NMDAR hypofunction. Altogether, our findings extend our current knowledge of reversal learning and behavioral flexibility beyond synaptic plasticity to the novel concepts of heterosynaptic LTD, repotentiation LTP, and flexible memory.

In conclusion, we have established that astrocytes play a crucial role in forming a flexible memory by enabling heterosynaptic LTD at unstimulated synapses, such that a new memory is easily formed when environment and circumstances change. These findings broaden our understanding of astrocytic roles in memory formation, and provide potential therapeutic targets for impaired cognitive flexibility in various psychiatric diseases.

## Acknowledgments

This study was supported by the Creative Research Initiative Program funded by National Research Foundation (NRF) of Korea (2015R1A3A2066619), Korea Institute of Science and Technology Institutional Program (project no. 2E26860), and Institute for Basic Science (IBS), Center for Cognition and Sociality (IBSR001-D2) to C.J.L. Thanks to Dr.Yuriy Pankratov for proofreading the manuscript.

## Author Contributions

W Koh, YE Chun, J Lee, MG Park, H Kang, J Woo, H Chun performed electrophysiological experiments. W Koh, M Park, HS Shim performed behavioral experiments. W Koh and S Kim performed slice imaging experiments. W Koh, MG Park, SJ Oh, S Lee, J Hong, J Feng performed molecular experiments. Y Li, H Ryu, J Cho, and CJ Lee gave technical support and conceptual advice. CJ Lee supervised the project. W Koh and CJ Lee wrote the manuscript.

## STAR★METHODS

### Contact for Reagent and Resource Sharing

Further information and requests for reagents may be directed to, and will be fulfilled by the corresponding author, C. Justin Lee.

### Animals

Mice were given ad libitum access to food and water and were kept under a 12:12-h light-dark cycle. All animals were housed in groups of 3–5 per cage. All animal care and handling was performed according to the directives of the Institutional Animal Care and Use Committee of Korea Institute of Science and Technology (Seoul, South Korea) and of Institute for Basic Science (Daejeon, South Korea).

### Stereotaxic virus injection into hippocampal CA1

Viruses used in this study were produced from Korea Institute of Science and Technology (KIST) Virus Facility (Seoul, South Korea) or Institute for Basic Science (IBS) virus facility (Daejeon, South Korea). Mice were anesthetized with isoflurane and mounted into stereotaxic frames (David Kopf Instruments, Tujunga, CA, USA). Viruses were bilaterally injected using syringe pump (KD Scientific, Holliston, MA, USA) into CA1 of hippocampus with the following coordinates (from bregma): anterior-posterior, −1.8 mm; medial-lateral, ±1.5 mm, dorsal-ventral, ±1.7 mm.

### Preparation of brain slice for the electrophysiological and imaging experiments

Brain slice were prepared as previously performed (Lee et al., 2007). Briefly, mice were anaesthetized with isoflurane and decapitated to isolate the brain. The brain was excised after decapitation, cut into 350-μm-thick transverse or coronal slices using vibrating microtome (DSK Linearslicer™ Pro7, DSK, Japan) in ice-cold, oxygenated (95% O_2_/5% CO_2_) sucrose-based dissection buffer containing 5 KCl, 1.23 NaH_2_PO_4_, 26 NaHCO_3_, 10 glucose, 0.5 CaCl_2_, 10 MgSO_4_, and 212.5 sucrose (in mM). Brain slices were left to recover for at least 1 hour before recording, and were used for whole-cell patch recording and imaging experiments.

For the fEPSP experiment, 400-μm-thick transverse of hippocampal slices were prepared and incubated in oxygenated artificial cerebrospinal fluid (aCSF) containing 124 NaCl, 5 KCl, 1.25 NaH_2_PO_4_, 2.5 CaCl_2_, 1.5 MgCl_2_, 26 NaHCO_3_ and 10 dextrose (in mM) at 28±1 °C for at least 1 hour.

### Whole-cell patch recording experiments

Whole-cell patch recordings were performed under the standard aCSF recording solution (130 NaCl, 24 NaHCO_3_, 3.5 KCl, 1.25 NaH_2_PO_4_, 1.5 CaCl_2_, 1.5 MgCl_2_ and 10 glucose (in mM)) saturated with 95% O_2_ and 5% CO_2_. Patch pipette electrodes (4-7MΩ) were fabricated from borosilicate glass (GC150F-10, Warner Instrument Corp., USA).

To measure exNMDAR current, recording electrodes were filled with an internal solution containing 135 CsMeSO_4_, 8 NaCl, 0.25 EGTA, 10 HEPES, 4 Mg-ATP, 0.3 Na_2_-GTP, and 5 QX314 (in mM) (pH adjusted to 7.3 with CsOH). Baseline current was stabilized under treatment of CNQX (20 μM), Bicuculline (10 μM), CGP 55845 (10 μM), and Strychnine (10 μM). The amplitude of exNMDAR current was measured by the baseline shift after 50 μM APV application. Signals were amplified using MultiClamp 700B (Molecular Devices, USA), and data was acquired and analyzed using a Digitizer 1550B (Molecular Devices, USA) and pClamp software (Molecular Devices, USA), respectively. To block Ca^2+^ signal in hippocampal astrocytes using BAPTA, patch electrodes were filled with an internal solution containing 123 CsCl, 1 MgSO_4_, 10 HEPES, 10 BAPTA, 100 Alexa fluor 488 hydrazide, 4 Mg-ATP, and 0.3 Na_2_-GTP (in mM) (pH adjusted to 7.35 with CsOH, and osmolality adjusted to 282 mOsmol/kg), as previously performed (refer). To measure evoked EPSC (eEPSC) and NMDA/AMPA, recording electrodes were filled with an internal solution containing 140 CsMeSO_4_, 8 NaCl, 1 MgCl_2_, 0.5 EGTA, 10 HEPES, 7 phosphocreatine di(tris) salt, 4 Mg-ATP, 0.3 Na_2_-GTP, and 5 QX314 (in mM) (pH adjusted to 7.3 with NMDG). Whole-cell voltage-clamp recordings were made from CA1 pyramidal neurons, and Schaffer collateral pathway was stimulated using a concentric bipolar electrode (CBBPE75, FHC, Bowdoin, ME, USA). AMPA-mediated current was recorded with holding at −60 mV and NMDA-mediated current was recorded with holding at +40 mV in the presence of CNQX. Stimulus intensity was adjusted to evoke an eEPSC of approximately 30–40% of the maximal amplitude.

To perform LTD and metaplasticity experiment, recording electrodes were filled with an internal solution containing 140 CsMeSO_4_, 8 NaCl, 1 MgCl_2_, 0.05 EGTA, 0.0244 CaCl_2_, 10 HEPES, 7 phosphocreatine di(tris) salt, 4 Mg-ATP, 0.3 Na_2_-GTP, 5 QX314 (in mM) (pH adjusted to 7.3 with NMDG).

In the whole-cell patch experiments, cells with a holding current lower than −100 pA or a change in the input resistance more than 30% were rejected.

### Ca^2+^ and NE imaging in brain slices

AAV-GFAP104-jRCaMP1a or AAV-GFAP104-GRAB_NE2m_ virus was injected, and coronal brain slices were prepared as described above. Imaging was acquired at 0.5 to 1 frame per second with a 60X water-immersion objective lens, and a 585-nm fluorescent imaging filter or 488-nm fluorescent imaging filter was utilized for jRCaMP1a or GRAB_NE2m_ imaging, respectively. Fluorescence Imaging was acquired with Imaging Workbench (Indec Biosystems), and analyzed with ImageJ software (NIH).

### Hippocampal fEPSP recording

Hippocampal slices were transferred to a submerged recording chamber and perfused with aCSF flowing at 2 mL/min. Schaffer collateral was stimulated using a concentric bipolar electrode (CBBPE75, FHC, Bowdoin, ME, USA), and fEPSP was recorded from stratum radiatum of CA1 using a glass pipette filled with aCSF (1-3 MΩ). Evoked fEPSP responses were amplified by an AC differential amplifier (DAM 80, World Precision Instruments, FL, USA) and digitized by BNC2110 (National Instruments). The slope of fEPSP response was analyzed by WinLTP v2.01 software (WinLTP Ltd., The University of Bristol, UK). The stimulation intensity was adjusted to obtain fEPSP slopes of 40-50 % to the maximum. During recordings, bath temperature was maintained at 28±1 °C by temperature controller (TC344B, Warner Instrument Corporation). Basal fEPSP response was monitored by electrical stimulations at 0.067Hz, and various stimulation protocols were delivered to test synaptic plasticity. High-frequency stimulation (HFS) consisted of 100 stimuli delivered at 100 Hz. 10 Hz stimulation consisted of 900 stimuli delivered at 10 Hz. Low-frequency stimulation (LFS) (1 Hz, 900 stimulations) consisted of 900 stimuli delivered at 1 Hz. For the simultaneous homosynaptic and heterosynaptic recordings, instead of concentric bipolar electrodes, borosilicate theta glass were fabricated and filled with aCSF to deliver focal stimulation on two independent pathways. Stimulation intensity was adjusted to acquire two independent pathways during paired-pulse ratio (PPR) test with 50 ms intervals, and the amplitude of each fEPSP was 0.1–0.4 mV.

### Primary astrocyte culture preparation

Primary astrocytes were prepared from P0-P3 of C57BL/6 mouse as described (Lee et al., 2007). Briefly, forebrain of mouse pup was dissected free of adherent meninges, minced and dissociated into single cell suspension by trituration. Cells were grown in Dulbecco’s modified Eagle’s medium (DMEM, Invitrogen) supplemented with 25 mM glucose, 10 % heat-inactivated horse serum, 10 % heat-inactivated fetal bovine serum, 2 mM glutamine and 1,000 units/ml penicillin–streptomycin. Cultured astrocytes were maintained at 37 °C in a humidified 5 % CO_2_ incubator. On the third day of culture (postnatal days 3, PND 3), cells were vigorously washed with repeated pipetting and the media was replaced to get rid of debris and other floating cell types.

The day after wash (PND 4), cells were re-plated onto cover-glass coated with 0.1mg/ml Poly D-Lysine (PDL), while various shRNAs were delivered to astrocytes by by electroporation. The electroporation was performed using the Microporator (Invitrogen) with an optimized voltage protocol (1200V, 2 pulses, 20 ms pulse width). Cell number for each transfection was around 2×10^6^. The following constructs were used; 5 mg pSicoR-SR-shRNA for SR knockdown experiment, 5 mg pSicoR-Best1-shRNA for Best1 knockdown experiment, and additional 5 mg pIRES2-mBest1-dsRED2 (shRNA insensitive form) or additional 5 mg pIRES2-mBest1-dsRED2 (W93C mutant) for control experiments.

### Sensor cell preparation

HEK293T cells were purchased from the Korean Cell Line Bank (Seoul National University) and cultured in DMEM (Invitrogen) supplemented with 10% fetal bovine serum (Invitrogen), 100 units/ml penicillin (Invitrogen), and 100 mg/ml streptomycin (Invitrogen) at 37 °C in a humidified 5% CO_2_ incubator. One day before the experiment for sniffer patch, HEK293T cells were transfected with 1:10 ratio of green fluorescence protein (pEGFP-N1) and pCINeo-NR1-1/NR2A(2D-S1) (5 mg per 60 mm dish) or 1:3 red fluorescence protein (pDsRed) and pCINeo-NR1-1/NR2A(2D-S1) using Effectene (Qiagen) following the manufacturer’s instructions. Additional 5 mM APV was supplemented in the medium to block the NMDA receptor-mediated cytotoxicity.

### Sniffer patch experiment

Sniffer patch was composed of Fura-2 imaging for Ca^2+^ from astrocytes and current recording from HEK293T cells expressing NR1-1/NR2A(2D-S1). On the day of experiment, the cover-slip in which astrocytes and sensor cells were seeded was incubated with 5 mM Fura-2 AM (mixed with 5 ml of 20% Pluronic acid) (Invitrogen) for 40 min, washed at room temperature, and subsequently transferred to the microscope stage for imaging. External solution contained (in mM): 150 NaCl, 10 HEPES, 3 KCl, 2 CaCl_2_, 2 MgCl_2_, 10 glucose and (pH adjusted to pH 7.3 and osmolarity adjusted to 325 mOsmol/kg). Intensity images of 510 nm wavelength were taken at 340 nm and 380 nm excitation wavelengths by using iXon EMCCD (ANDOR). To induce astrocytic Ca^2+^, 500 μM TFLLR was applied with pressure (20 lbf in–2, 100ms) using Picospritzer (Parker Instrument, USA). Two resulting images were used for 340/380 ratio calculation in Imaging Workbench version 6.2 (Indec Systems). NR1-1A/NR2A(2D-S1)-mediated currents were recorded from HEK293T cells under voltage clamp (V_h_ = –70mV) using Axopatch 200A amplifier acquired with pClamp 10.4 (Molecular Devices). Recording electrodes (4–7MΩ) were filled with (mM): 110 Cs-Gluconate, 30 CsCl, 0.5 CaCl_2_, 10 HEPES, 4 Mg-ATP, 0.3 Na_2_-GTP, and 10 BAPTA (pH adjusted to 7.3 with CsOH and osmolarity adjusted to 290–310 mOsm/kg with sucrose). For simultaneous recording, Imaging Workbench was synchronized with pClamp 10.4.

### Two-cell assay

To investigate co-release of D-serine and glutamate via Best1, two-cell assay was performed. Preparation of sensor cell was performed as described above. For source cell preparation, HEK cells were transfected with pIRES2-mBest1-dsRED2 or pIRES2-mBest1(W93C)-dsRED2 (7 mg per 60 mm dish) using Effectene (Qiagen). The internal solutions for source cell contained (in mM); glutamate without D-serine, 90 CsCl, 50 glutamate, 10 HEPES, 5 (Ca^2+^)-EGTA-NMDG, 2 MgCl_2_, 0.3 Na_2_-GTP, 4 Mg-ATP (pH 7.3 with CsOH, 289 mOsmol by adding sucrose); glutamate with D-serine, 40 CsCl, 50 glutamate, 50 D-serine, 10 HEPES, 5 (Ca^2 +^)-EGTA-NMDG, 2 MgCl_2_, 0.3 Na_2_-GTP, 4 Mg-ATP (pH 7.3 with CsOH, 289 mOsmol by adding sucrose). A pair of one sensor cell and one source cell were patched, and the responsive current from sensor cells was measured under voltage clamp (V_h_ = –70mV), while the source cell is ruptured. Sensor current was measured as described above.

### Permeability assay

To estimate the D-serine permeability of Best1, Best1 current was measured from Best1-expressing HEK293T cell as previously performed (Lee et al., 2010), with various concentration of substitution for chloride to D-serine. The internal solution contained (in mM); 100 CsCl, 20 tetraethylammonium (TEA)-Cl, 8.7 CaCl_2_, 10 HEPES, 10 BAPTA, 3 Mg-ATP, 0.2 Na_2_-GTP, and 0.5 MgCl_2_, (pH was adjusted to 7.2 with CsOH); when D-serine was included, it replaced an equimolar amount of CsCl. Osmolarity was adjusted to 287 mOsmol by adding sucrose. For these experiments, the external solution contained (in mM); 126 NaCl, 10 HEPES, 20 glucose, 1.8 CaCl_2_, 1.2 MgCl_2_, and 10 TEA-Cl (pH 7.4 with NaOH).

### Immunohistochemistry

Adult mice were deeply anesthetized with 2% avertin and perfused with 0.1 M PBS followed by 4% paraformaldehyde. Brains were postfixed in 4% paraformaldehyde at 4°C for 24 hours and 30% sucrose at 4°C for 48 hours. Frozen brains were then cut into 30 μm coronal cryosections. Sections were blocked in 0.1 M PBS containing 0.3% Triton X-100 (Sigma) and 2% Donkey Serum (GeneTex) for 30 min at room temperature. Primary antibody was applied at the appropriate dilution and incubated overnight at 4°C. Incubated sections were washed three times with 0.1 M PBS and incubated in secondary antibody for two hours. After three rinses in 0.1 M PBS and DAPI staining at 1:1000 (Pierce), the sections were mounted on polysine microscopic glass slides (Thermo Scientific). Images were scanned with the Axio Scan Z1 automated slide scanner (Zeiss) using ZEN 2 (blue edition) slidescan software (Zeiss).

### Western blotting

Western blotting was performed as previous (Woo et al., 2012). Briefly, to test the expression of NMDAR1, hippocampi were dissected from WT and Best1 KO, and the tissues were lysed with RIPA buffer (50 mM Tris-HCl, pH 7.4, 150 mM NaCl, 5 mM EDTA, 1 mM PMSF, and 1% NP-40) containing a protease-inhibitor cocktail. Obtained protein lysates were separated by SDS-PAGE using 10% gels and blotted onto PVDF membranes. The blots were incubated with rabbit anti-NMDAR1 (ab109182, Abcam) and anti-β-actin antibody (1:2,000; ab133626, Abcam). To test knockdown efficiency of SR, SR shRNA was transfected as described above, and protein lysates were acquired three days after transfection. The blots were incubated with mouse anti-SR (sc-365217, Santa Cruz) and anti-β-actin antibody. Appropriate horseradish peroxidase-conjugated secondary antibodies (Jackson ImmunoResearch) were used for detection by enhanced chemiluminescence (GE Healthcare). The band intensity was acquired by ImageQuant LAS 4000 (GE Healthcare) and quantified using ImageJ software (NIH).

### Behavioral tests

#### Contextual fear conditioning

Contextual fear conditioning test was performed as previously described (Jung et al., 2016). The chamber with a stainless-steel floor was located in a sound-proof box with a camera mounted on its ceiling (Med associates, Inc., St. Albans, VT, USA). On the first day, mice were allowed to explore the chamber freely for 3 min, and received six foot shocks separated by 1 min (0.4 mA, 2 seconds). Twenty-four hours after the training, the mice were placed in the chamber for 12 min and their behavioral responses were videotaped. Freezing response, defined as an absence of any movement except breathing for >1 s, was measured and divided by total time to.

#### Passive avoidance test

Passive avoidance test were performed as previously described (Jo et al., 2014). Briefly, mouse was placed in the two-chamber (light and dark) box with a constant current shock generator (MED Associates). On the acquisition, mouse was released into light chamber for one minute (habituation), and a gate between two chambers was opened. When mouse passed the gate, the gate immediately is closed, and aversive electric shock (0.5 mA, 2 seconds) was delivered through the floor. After the delivery of electric shock, mouse was returned to the home cage. Retention test was performed 24 hours after acquisition test. For retention test, mouse was placed in the light chamber and the door opened 1 minute later, and the latency of entry to dark chamber was recorded.

#### Morris water maze

Morris water maze experiments were performed as previously described (Park et al., 2015). Briefly, for hidden platform Morris water maze experiments, animals were trained to find a hidden platform (10 cm diameter, 1 cm under the water surface) placed in a fixed location in a water maze (1.2 m diameter) filled with water (25 °C) made opaque by the addition of nontoxic white paint (Weather tough Forte, Bristol Paints). The water maze was surrounded by a black circular curtain (placed 70 cm away) that held 3 salient visual cues. The releasing point was randomly distributed across 4 quadrants of the pool and the animal was allowed maximum 60 sec to find the hidden platform. If escape did not occur within 60 sec, the animal was manually guided to the platform where they stayed on for 30 sec. The training consisted of 4 trials/day (10 min inter-trial interval, ITI) for 7 days. On training days 4 and 8, animals were given 60 sec probe tests to test their spatial memory. After 7 days of acquisition, the hidden platform was placed on the opposite quadrant to test spatial reversal learning for 3 additional days and the final probe test. For the D-serine application, D-serine (600 mg/kg) was injected intraperitoneally 20 min before the first trial of each day during acquisition session. Control group consisted of half saline-injected mice and half naive mice. The amount of saline injection was set equal to the amount of saline in which D-serine was dissolved. Neither D-serine nor saline was injected during the following reversal learning session.

For the visible platform test, animals were trained to find a visible platform (10 cm diameter, 1 cm above the water surface) marked with a salient black tape for 2 days (4 trials/day, 10 min ITI). If the animal found the platform, the animal remained on the platform for 30 sec. During the test session after acquisition (day 3, trial 9), the platform was moved to a new location (adjacent right quadrant). And the animals were released in the pool equidistant from the original and new location. An automated tracking system (Noldus, Netherlands) was used to monitor and analyze the number of platform crossing, and the amount of time spent in each of the four quadrants.

For the Best1 rescue experiments, experimental conditions were set as above, except the size of water maze (1.5 m diameter) and the number of trials (3 trials/day, 10 min ITI).

## Statistical analysis

Statistical analyses were performed using Prism 9. All data were presented as mean ± SEM. No statistical method was used to predetermine sample size. Sample sizes were empirically determined based on our previous experiences or other literatures. Experimental groups were balanced in terms of animal age, sex and weight. Animals were genotyped prior to the experiment, and they were treated in the same way. Animals were randomly and evenly allocated to each experimental condition. To perform the group allocation in a blind manner during data collection, animal preparations and experiments were carried out by different researchers. Statistical significance was set as *p < 0.05, **p < 0.01, ***p < 0.001.

## Supplemental Information

**Figure S1.**
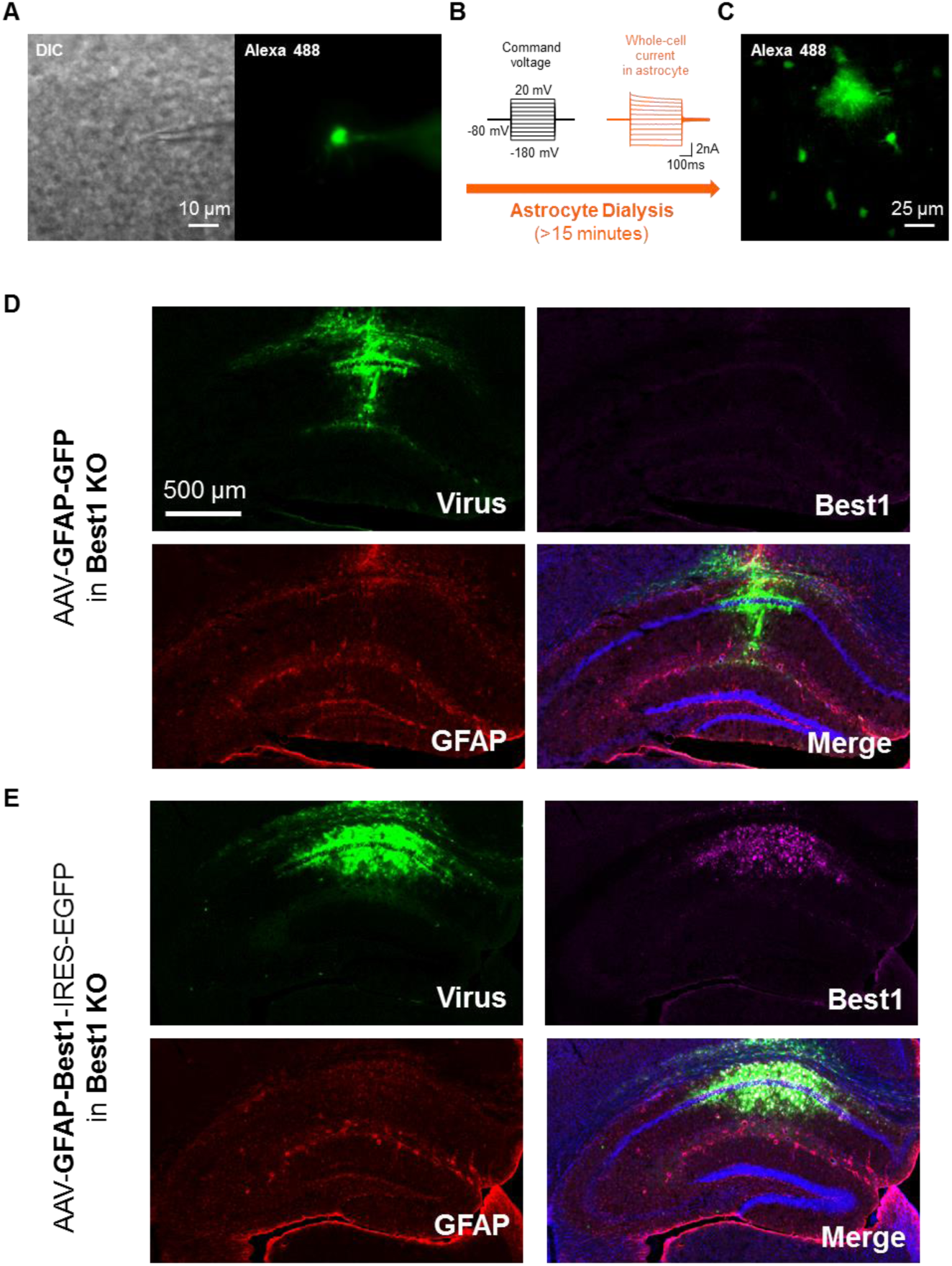
Astrocytic dialysis through patch-clamp recording and rescue of astrocytic Best1 in hippocampal CA1 with astrocyte-specific AAV viruses. Related to Figures 1, 3, 7. (A) DIC and fluorescence image of hippocampal astrocyte in CA1 stratum radiatum after whole-cell patch recording with BAPTA, Alexa 488 fluorophore-containing internal patch pipette. (B) Passive conductance from astrocytes during astrocytic dialysis for more than 15 minutes. (C) Representative images of astrocytic dialysis with Alexa 488 fluorophore. (D) AAV-GFAP-GFP (control virus) injection into hippocampal CA1 of Best1 KO. Virus-infected cells (green) show no Best1 expression (magenta). (E) AAV-GFAP-Best1-IRES-EGFP (rescue virus) injection into hippocampal CA1 of Best1 KO. Virus-infected cells (green) show Best1 expression (magenta).

**Figure S2.**
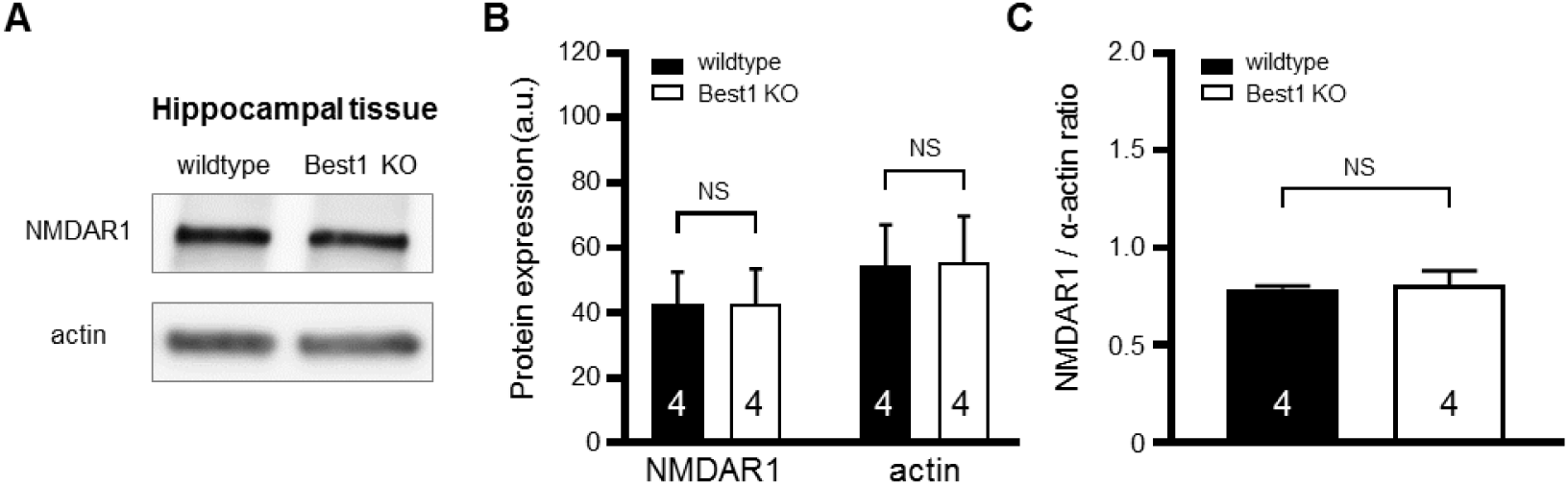
Hippocampal NMDAR1 expression in WT and Best1 KO. Related to Figures 1. (A) Western blot results for NMDAR1 and actin from hippocampal tissues of WT and Best1 KO. (B) Quantification of the western blot results from hippocampal tissues of WT and Best1 KO. Number indicates animal number of each condition. (C) Ratio of NMDAR1 / actin expression in hippocampi of wildtype and Best1 KO. Number indicates animal number of each condition. Data are presented as mean ± SEM. NS > 0.05.

**Figure S3.**
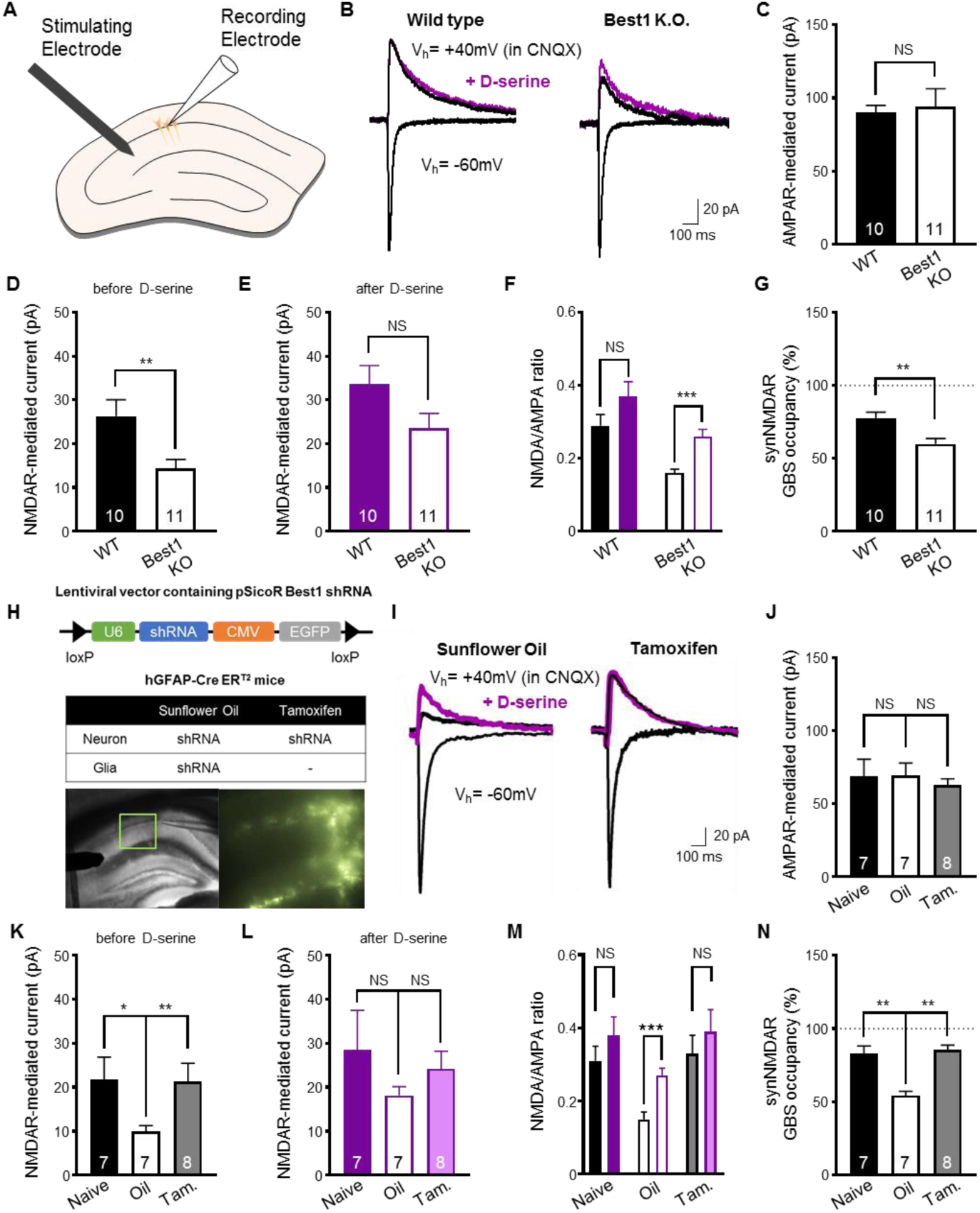
SynNMDAR in WT and Best1 KO. Related to Figures 1. (A) Scheme of synNMDAR recording. (B) Representative AMPAR-, NMDAR-mediated current, and NMDAR-mediated current with 100 µM D-serine application in WT and Best1 KO. (C-G) Summary graph of AMPAR (C), NMDAR (D), NMDAR after 100 µM D-serine application (E), NMDAR/AMPAR ratio (F), and estimated synNMDAR glycine modulatory site (GMS) occupancy (%) (G) in WT and Best1 KO. (H) Cell-type specific manipulation of Best1 shRNA in hGFAP-CreERT2 mice. General knockdown of Best1 with sunflower oil (Oil, control), and astrocyte-specific rescue of Best1 with tamoxifen (Tam.) application. (I) Representative AMPAR-, NMDAR-mediated current, and NMDAR-mediated current with 100 µM D-serine application from Oil- and Tam.-injected group. (J-I) Summary graph of AMPAR (J), NMDAR (K), NMDAR after 100 µM D-serine application (L), NMDAR/AMPAR ratio (M), and estimated synaptic NMDAR glycine modulatory site (GMS) occupancy (%) (N) in naïve, Oil- and Tam.-injected group. Data are presented as mean ± SEM. *p < 0.05; **p < 0.01; ***p < 0.001.

**Figure S4.**
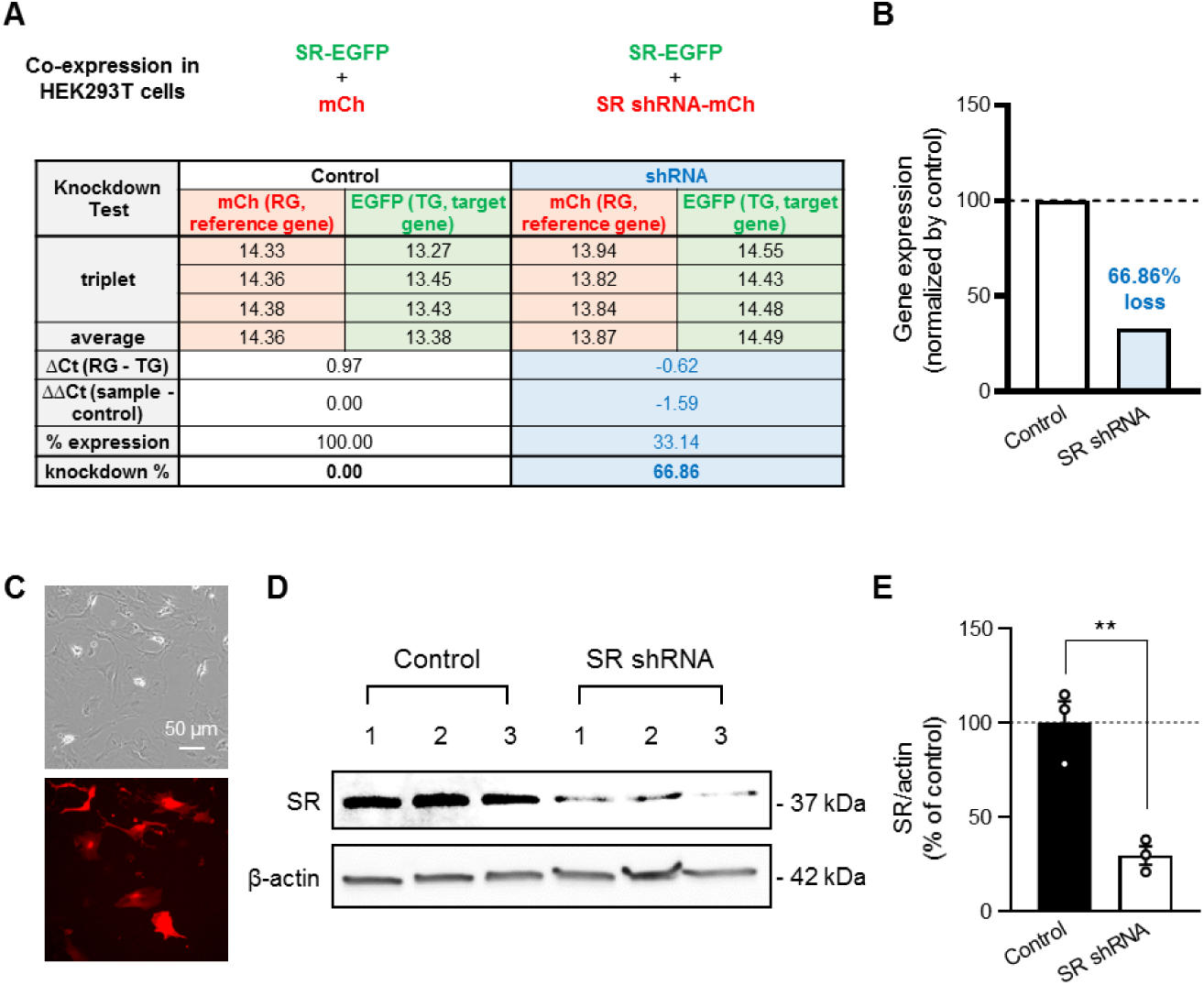
Validation of SR shRNA. Related to Figures 2 and 3. (A) Upper: SR shRNA efficiency was accessed by co-expression of SR-EGFP and shRNA (or control vector) in HEK293T cells. Lower: Results of Quantitative RT-PCR and analyzed results by ΔΔCt method. (B) Summarized results of mRNA knockdown efficiency of mSR shRNA. (C) Transfection of SR shRNA in primary astrocytes. (D) Western blot results for SR in control and SR shRNA conditions. Numbers indicate different culture batches. (E) Ratio of SR / actin expression. Dots indicate different culture batches. Data are presented as mean ± SEM. *p < 0.05; **p < 0.01; ***p < 0.001.

**Figure S5.**
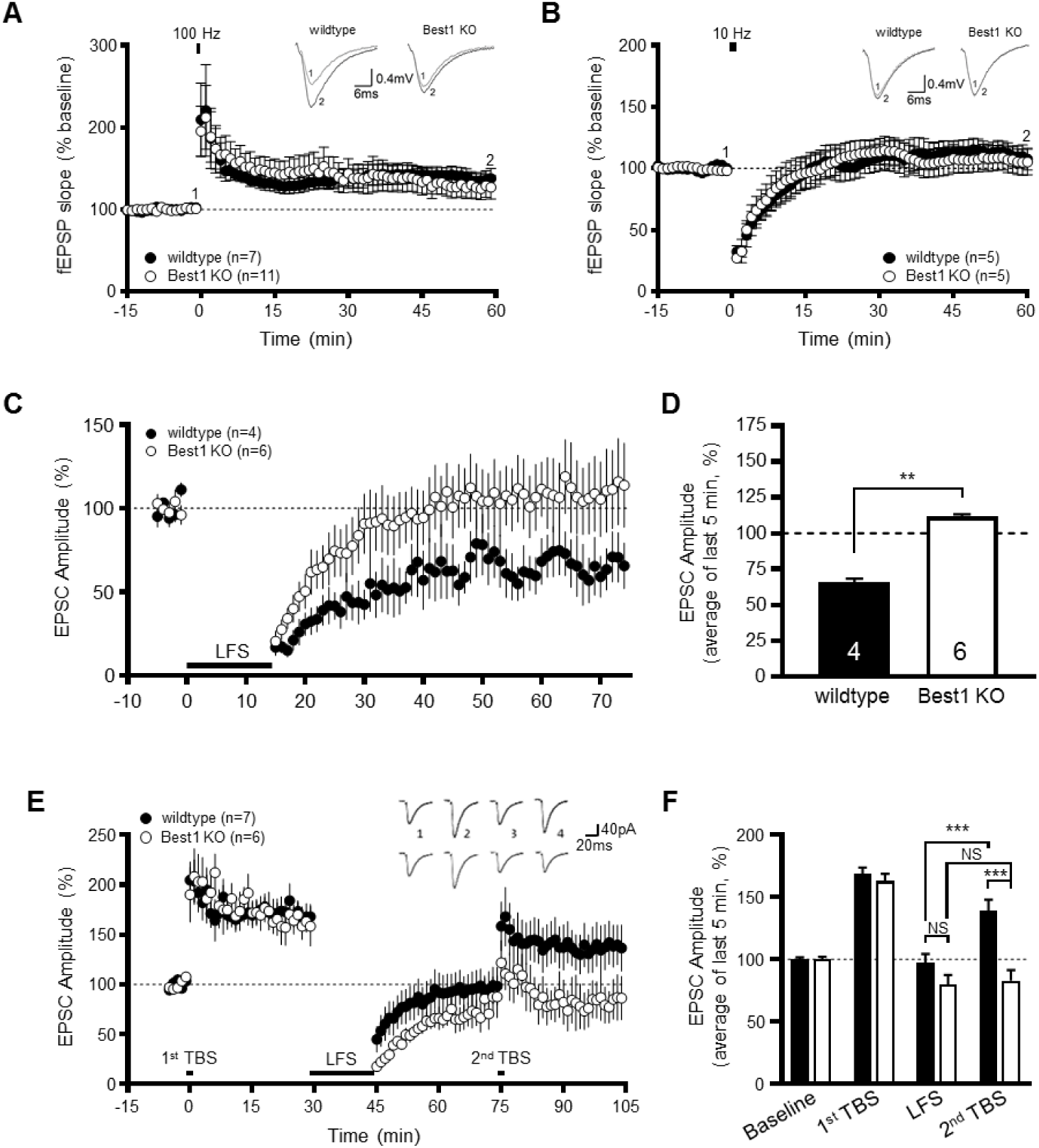
Hippocampal synaptic plasticity in Best1 KO. Related to Figures 3 and 6. (A) HFS (100 stimuli at 100 Hz)-induced LTP in WT and Best1 KO under fEPSP recording. (B) 10 Hz stimulation (900 stimuli) in WT and Best1 KO under fEPSP recording. (C) LFS (900 stimuli at 1 Hz)-induced LTD in WT and Best1 KO under excitatory postsynaptic current (EPSC) recording from CA1 pyramidal neuron with whole-cell patch-clamp. (D) Summary graph of LTD induction in WT and Best1 KO under EPSC recording. (E and F) Time course of the normalized EPSC changes (E) and summary graph (F) of 1^st^ TBS (theta-burst stimulation)-induced potentiation, LFS-induced depotentiation, and 2^nd^ TBS-induced repotentiation in WT and Best1 KO. Data are presented as mean ± SEM.. *p < 0.05; **p < 0.01; ***p < 0.001.

**Figure S6.**
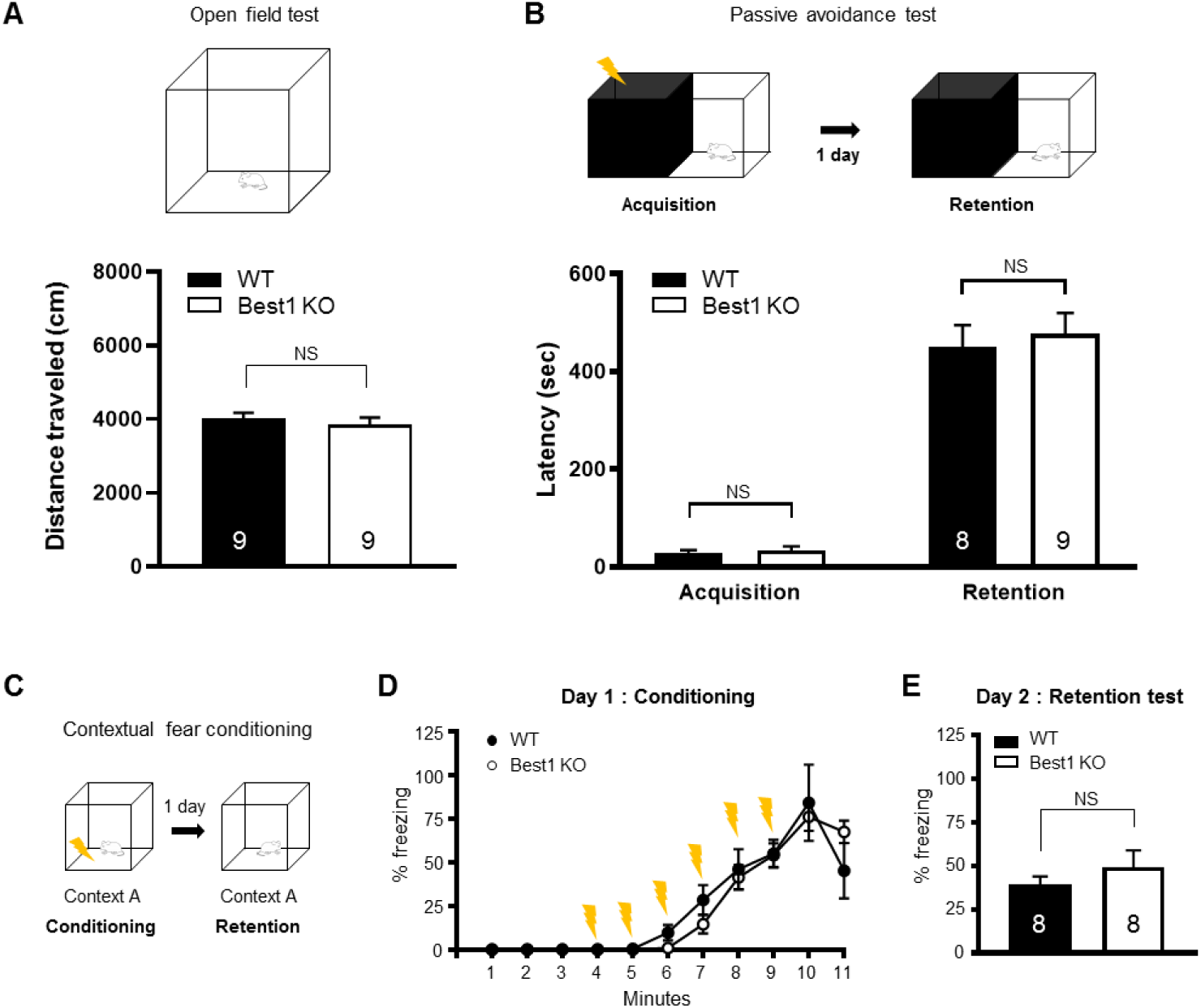
Best1 KO mice have impaired reversal learning, but not other hippocampus-dependent spatial learning. Related to Figure 7. (A) Top, Open field test (OFT). Bottom, Summary graph of distance traveled during OFT. (B) Top, Passive avoidance test (PAT). Bottom, Summary graph of latency to the dark room before and after shock-association in WT and Best1 KO. (C) Contextual fear conditioning (CFC). Retention test was performed in the same context, the day after fear conditioning with electric shock (yellow symbol). (D) Percentage of freezing time during conditioning session in WT (black) and Best1 KO (white). Yellow symbols represent the delivery of electric shock. (E) Percentage of freezing time in retention session the day after conditioning. Numbers indicate the number of animals. Data are presented as mean ± SEM. *p < 0.05; **p < 0.01; ***p < 0.001.

**Figure S7.**
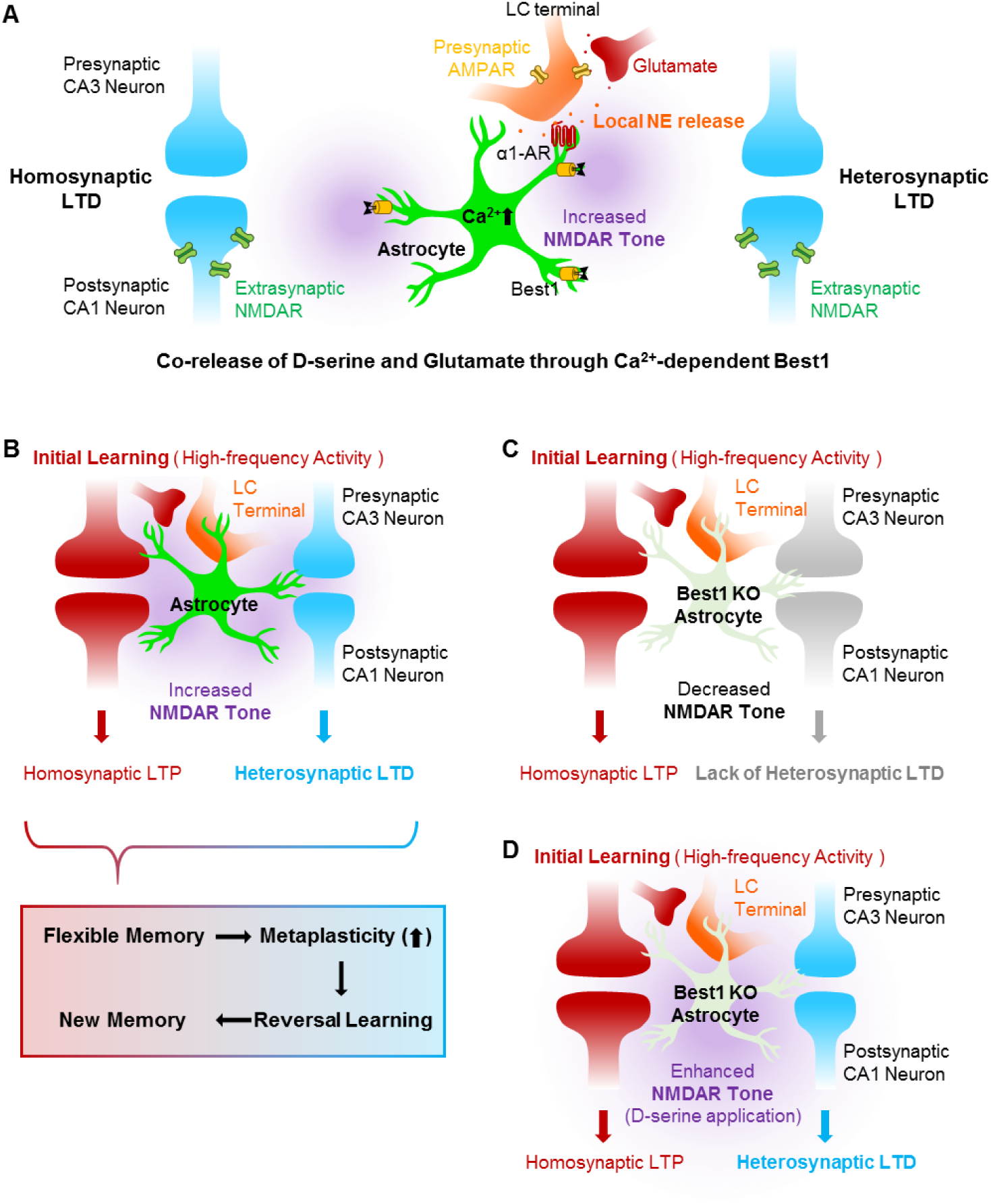
Schematic illustration of a working model for Best1 in flexible memory. Related to Figure 7. (A) NMDAR tone is regulated by astrocytic co-release of D-serine and Glutamate through Ca^2+^-dependent Best1. (B) Local NE release activates astrocytes, increases NMDAR tone, and induces heterosynaptic LTD. Heterosynaptic LTD enable further plasticity (metaplasticity) and reversal learning. (C) Heterosynaptic LTD is impaired in Best1 KO. (D) The impaired heterosynaptic LTD is restored by D-serine application.

